# Comparative Analysis of Single-Nucleus and Single-Cell RNA Sequencing in Human Bone Marrow Mononuclear Cells: Methodological Insights and Trade-offs

**DOI:** 10.1101/2025.09.08.675012

**Authors:** Reza Ghamsari, Carolyn A. de Graaf, Rachel Thijssen, Yupei You, Nigel H. Lovell, Hamid Alinejad-Rokny, Matthew E. Ritchie

## Abstract

Bone marrow mononuclear cells (BMMCs) are a heterogeneous pool of hematopoietic progenitors and mature immune cells that collectively sustain hematopoiesis and coordinate immune responses. The bone marrow serves not only as the primary site for blood cell production but also as a niche for various disorders, including blood cancers. Advances in single-cell RNA sequencing (scRNA-seq) and single-nucleus RNA sequencing (snRNA-seq) have significantly enhanced our understanding of the cellular biology and molecular dynamics within this complex microenvironment. The choice between these two approaches, however, is often shaped or constrained by the study design, such as research objectives, sample type, and preservation conditions. Consequently, methodological differences in library preparation and transcript capture efficiency can introduce systematic biases that complicate downstream analyses and interpretation, underscoring the need to identify and account for method-specific features. In this study, we conducted a comparative analysis of matched snRNA-seq and scRNA-seq datasets from 11 pairs of healthy donor bone marrow mononuclear cell samples, generated using the popular 10x Genomics platform. We evaluated method-specific biases using multiple quality metrics and compared cell type proportions and transcriptomic signatures captured by each approach. Integrative analysis of these datasets is feasible but not advisable due to systematic gene length biases that were observed between these approaches. Our results showed that despite inherent differences in library complexity, both protocols reliably captured all major cell types. This comparative analysis highlights intrinsic differences between snRNA-seq and scRNA-seq data, providing valuable insights into their respective advantages, limitations, and trade-offs. These findings can assist researchers in selecting the optimal method tailored to specific biological questions and sample characteristics, and also enable more method-aware data analysis and interpretation.

## Introduction

Bone marrow mononuclear cells (BMMCs) are a heterogeneous pool of hematopoietic cells, essential for maintaining blood cell production and immune function. Studying the complex and heterogeneous microenvironment of bone marrow at the single cell level is essential for advancing our understanding of hematopoiesis, immune regulation, and the pathogenesis of various blood disorders (Ennis et al., 2022; Zhao et al., 2012). Over the past decade, advances in sequencing technologies have enabled high-throughput transcriptome profiling at single-cell resolution, significantly deepening our understanding of hematopoiesis (Lee et al., 2023b). In particular, microfluidic droplet-based methods have emerged as the leading technology for high-throughput singlecell transcriptomic profiling (Tirosh and Suva, 2024). Among the most widely adopted of these are singlecell RNA sequencing (scRNA-seq) and single-nucleus RNA sequencing (snRNA-seq), commercialised by 10x Genomics (Woo and Eyun, 2025). Depending on sample availability, time constraints, and the specific biological questions, both approaches are amenable to BMMC transcriptome profiling. snRNA-seq facilitates the use of frozen clinical samples, alleviating time constraints associated with fresh tissue processing, and allows multiplexing, which enables the analysis of multiple samples simultaneously to reduce batch variation (Slyper et al., 2020; Van Melkebeke et al., 2024). Compared to scRNA-seq, snRNA-seq is more able to capture newly transcribed RNA, and thus early transcriptional responses (Meyer et al., 2023); however, these timing discrepancies are far smaller than sampling resolution, as most mRNA is exported from the nucleus in under a second (Kubitscheck and Siebrasse, 2017). Despite the ability of both methods to capture the transcriptome profiles of thousands of cells, evaluating their performance in different tissues is important (Deleersnijder et al., 2021). Therefore, further cross-validation of single-cell methodologies is necessary to confirm the reliability and reproducibility of these findings across diverse samples and conditions (Oetjen et al., 2018).

While scRNA-seq involves isolating intact, viable cells to capture the transcriptomic landscape of both cytoplasmic and nuclear RNA, snRNA-seq relies on the isolation of nuclei, thus capturing only nuclear RNA. Notably, snRNA-seq was primarily developed for cells that are difficult to dissociate, such as neurons and frozen tissues with damaged cell membranes (Kim et al., 2023). Both approaches require different degrees of tissue dissociation to generate high-quality single-cell or singlenucleus suspensions prior to RNA extraction (Deleersnijder et al., 2021; Kersey et al., 2025). The tissue dissociation step is critical yet prone to complications, as harsher dissociation conditions may damage fragile cells, compromising their viability and leading to the loss of certain cell types. Conversely, milder dissociation conditions might result in incomplete dissociation, leaving extracellular matrix-embedded cells underrepresented. Additionally, a higher number of cells is needed to capture rare cell types (Deleersnijder et al., 2021). Previous research has demonstrated that including intronic reads during preprocessing improves the concordance between transcriptomic profiles obtained from scRNA-seq and snRNA-seq (Chamberlin et al., 2024). This is consistent with the nature of snRNA-seq, which captures nuclear RNA enriched in unspliced pre-mRNA, resulting in a greater intronic read content compared to scRNA-seq (Truong et al., 2023; Habib et al., 2017; Kim et al., 2023). These fundamental methodological differences, along with distinct laboratory workflows, have raised ongoing questions among researchers regarding potential biases and discrepancies between the two approaches. For example, the ability to recover different cell types is a critical consideration when benchmarking single-cell methodologies (Ding et al., 2020). In two separate studies comparing cell type compositions with histological annotations in rabbit retina and human normal lung tissue, snRNA-seq demonstrated a more accurate reflection of cell type composition (Santiago et al., 2023; Renaut et al., 2024). These findings support the expectation that nuclear profiling is less cell-type biased compared to scRNA-seq (Bakken et al., 2018).

Beyond pure transcriptomics, the ability to measure multiple molecular modalities within a cell simultaneously has significantly enhanced our capability to resolve heterogeneous cellular populations (Guo et al., 2020). Multimodal approaches can reveal new relationships and interactions that occur between different molecular layers and ultimately incorporate these relationships to disentangle causal events in gene regulation and cellular phenotypes (Ma et al., 2020; Argelaguet et al., 2021). The isolation of nuclei in snRNA-seq facilitates access for the transposase enzyme to chromatin within the nucleus, enabling the simultaneous profiling of chromatin accessibility (scATAC-seq) and gene expression. This capability is harnessed by the 10x Genomics Chromium Single Cell Multiome ATAC + Gene Expression kit, which allows paired measurement of the transcriptome and chromatin accessibility from the same nucleus, providing insights into gene regulatory mechanisms at single-cell resolution (10X Genomics, a). In contrast, the presence of an intact cellular membrane in scRNA-seq is essential for compatibility with techniques that require surface protein detection. This feature enables the application of methods like Cellular Indexing of Transcriptomes and Epitopes by Sequencing (CITE-seq), which utilises 10x Genomics’ Feature Barcoding technology. This technology allows for the simultaneous measurement of gene expression and cell surface proteins at the single-cell level using oligonucleotide-labelled antibodies. CITE-seq provides an additional layer of phenotypic information, facilitating more precise cell type identification and functional characterisation (Stoeckius et al., 2017). Additionally, scRNA-seq and snRNA-seq can effectively serve as bridges between unpaired modalities, such as chromatin accessibility and protein profiling. For instance, better cell type annotation using CITE-seq can be linked to ATAC-seq data through RNA profiles, facilitating comprehensive multi-omics analyses at the single-cell level (Lin et al., 2022; Hao et al., 2021).

As both scRNA-seq and snRNA-seq approaches gain widespread adoption, it becomes increasingly important to assess the efficiency, capabilities, and potential biases of each method in capturing the transcriptomic profile of individual cells. Relatively few studies have performed paired-sample comparative analyses of snRNA-seq and scRNA-seq, and none to date have involved BMMCs. One example is a recent publication involving paired samples from human bladder tissue; however, this work was limited to a single human donor, with only one replicate per region, all processed within a single laboratory (Santo et al., 2025). In this study, we conducted a comparison of scRNA-seq and snRNA-seq datasets derived from different healthy human donors and processed across different laboratories to evaluate the consistency and complementarity of these two approaches in studying BMMCs. By highlighting the biases inherent in the transcriptomic profiles captured by each method, we aimed to ascertain whether the primary transcriptomic signatures identified by scRNA-seq are preserved within snRNA-seq data, helping researchers to select the most appropriate method based on their specific research objectives and sample types.

## Results

We conducted a comparative analysis of publicly available datasets comprising 11 matched pairs of bone marrow mononuclear cell samples from healthy donors (Figure 1A1-3), generated using scRNA-seq and snRNA-seq approaches across four different laboratories (Luecken et al., 2021). We evaluated these two methodologies using various quality indices to better understand different aspects of the data and to assess their respective capacities for capturing transcriptomic signal, as well as their ability to distinguish between different cell populations (Figure 1B1-11).

**Figure 1.**
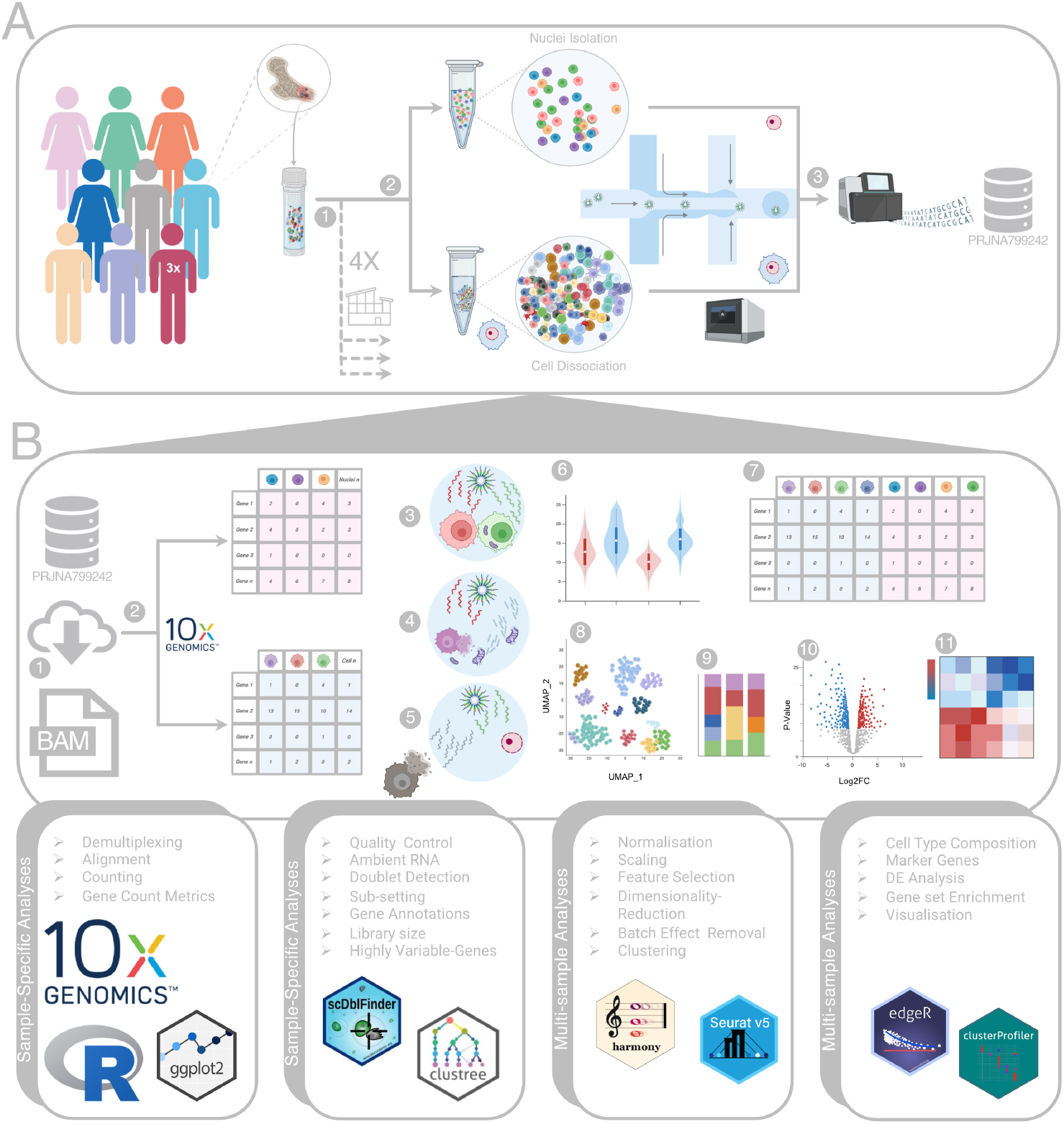
Schematic representation of the study design, dataset and analysis workflow. **A)** This study re-analysed a publicly available dataset comprising 11 matched pairs of BMMCs samples from healthy donors. These samples were processed across four (A1) different laboratories, where each donor’s sample was divided into two aliquots: one processed using scRNA-seq and the other using snRNA-seq (A2). Libraries for both scRNA-seq and snRNA-seq were sequenced on the Illumina NextSeq 2000 platform (A3). **B)** Comparative analyses between the two approaches were conducted using the Seurat pipeline and other R packages. For sample-specific analyses, the raw data were downloaded (B1) and aligned to the reference genome using 10x Genomics’ Cell Ranger software (B2). The QC steps include doublet detection (B3), removing cells with high mitochondrial gene percentage (B4), ambient RNA removal (B5) and a biology-aware QC approach (B6), which was performed first on each sample separately. Multi-sample analyses, followed by merging different samples (B7), applying batch effect removal, clustering (B8), cell type annotation (B9), differential expression (DE) analysis (B10) and subsequent downstream analyses (B11). Figure created with BioRender.com.

### Sample-specific Analyses

Conducting quality control (QC) on a per-sample basis enables filtering that accounts for sample-specific variability, ensuring the retention of high-quality cells unique to each dataset. A total of 211,590 filtered barcodes were recovered across all samples using Cell Ranger, and the resulting count matrices were used for downstream analyses. Comparisons of various attributes from Cell Ranger’s summary reports and BioMart annotations revealed that scRNA-seq samples exhibited a significantly higher number of genes detected per cell and total number of reads, along with lower sequencing saturation, compared to snRNA-seq samples (Figure 2A-C). This reflects greater transcriptomic coverage and library complexity inherent to scRNA-seq. Additional attribute comparisons are presented in Supplementary Figure S1-S3. Notably, examining the correlation between the number of genes and the number of counts per cell, erythroblasts in the scRNA-seq data displayed a distinct trend, characterised by larger library size concentrated in a smaller number of genes (Figure 2D, E). Comparing signals across different genomic intervals, snRNA-seq samples exhibited a higher proportion of reads mapping confidently to intronic regions (Figure 2F). For gene-level annotations, snRNA-seq data exhibited a higher representation of longer genes, whereas scRNA-seq data were enriched for shorter genes (Figure 2G). To ensure that these biases were not driven by the overexpression of a few highly expressed genes, we removed several such genes, including ribosomal (RB) and hemoglobin (HB) genes. While this eliminated some prominent peaks in the length distribution plots, it did not alter the overall gene-length bias trend (Supplementary Figure S4A,B). To further assess this bias, we analysed an independent dataset comprising a pool of cells from eight distinct cancer cell lines generated in our laboratory using both sequencing methods, and observed a similar gene length bias between the two methods (Supplementary Figure S4C). Comparison of different gene biotype annotations revealed significant composition differences between the two methods. However, the presence of a gene-length bias introduces a confounding factor that complicates the interpretation of these results. We observed that scRNA-seq samples generally exhibited higher mitochondrial (MT) gene proportions than snRNA-seq (Figure 2H), as expected given the lack of cytoplasmic content in the latter. Interestingly, some snRNA-seq samples exhibited a bimodal distribution of MT gene expression (Supplementary Figure S5), potentially indicating the presence of damaged cells or partially isolated nuclei. snRNA-seq data is also significantly more susceptible to ambient RNA contamination compared to scRNA-seq (Figure 2I). Our findings did not show a notable difference in doublet rates between scRNA-seq and snRNA-seq. However, we observed a significant positive correlation between the number of cells captured and the doublet rate (Figure 2J).

**Figure 2.**
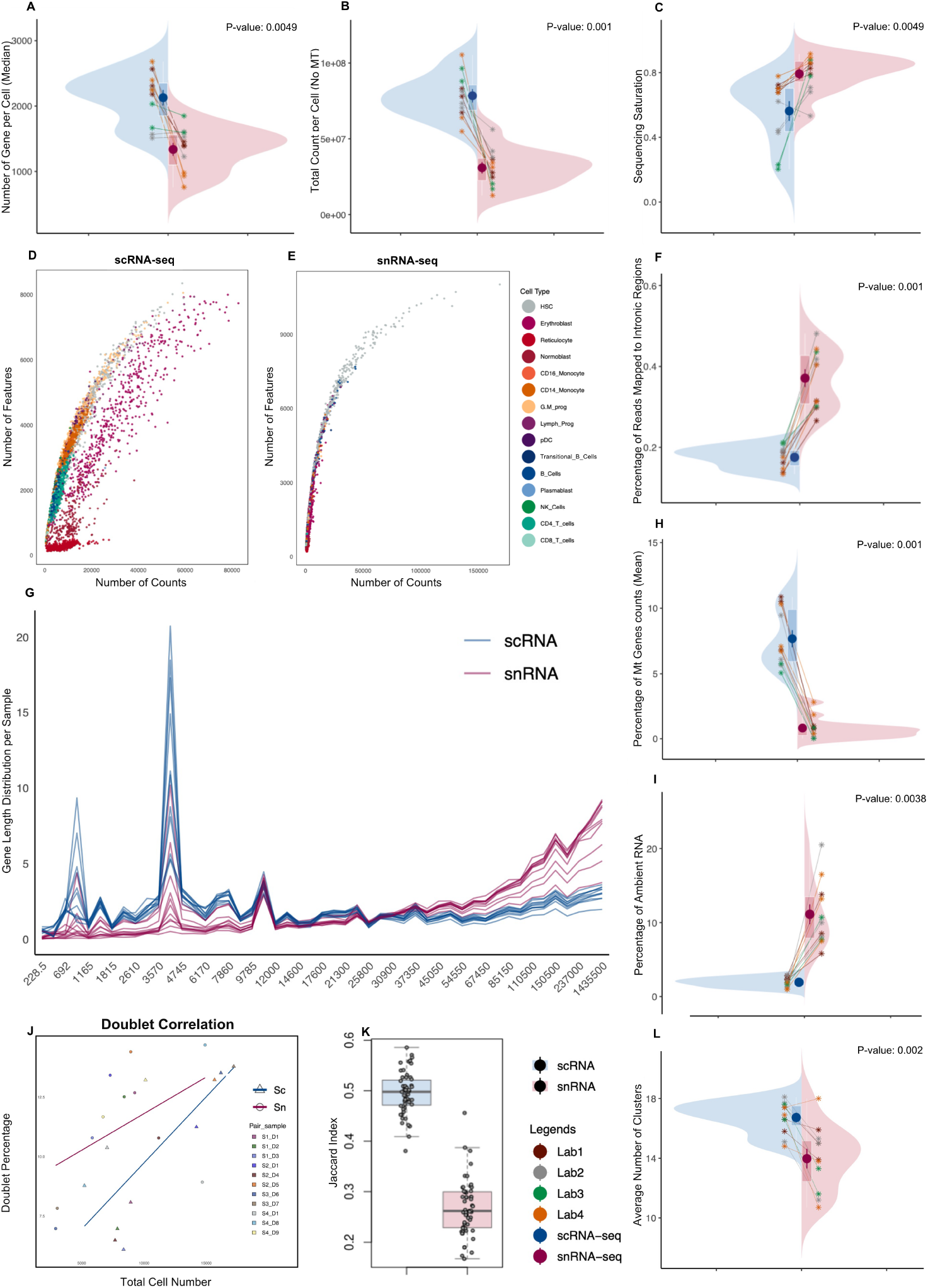
Sample-level comparisons between scRNA-seq and snRNA-seq. **A–C)** Split violin plots display the distribution of key sequencing and cell quality metrics, with red representing snRNA-seq and blue representing scRNA-seq. Each pair of samples is connected by a line. Samples are coloured by the laboratory to indicate batch origin (A, B, C, F, H, I, L). Median number of genes detected per cell for each sample (A), Library size per sample after removing MT genes (B) and Sequencing saturation (C). **D–E)** Scatter plots showing an example of the correlation between library size and number of genes per cell in scRNA-seq (D) and snRNA-seq (E), coloured by cell type. Scatter plots for all samples are shown in Figure S7. **F)** Split violin plots showing proportion of reads mapped to intronic regions per sample. **G)** Distribution of total UMI counts per gene length bin across 50 gene length bins across all 22 samples. **H–I)** Split violin plots comparing the percentage of MT gene content (H) and ambient RNA (I). **J)** Correlation between the percentage of detected doublets and the total number of cells recovered per sample. **K)** Boxplots showing Jaccard scores representing similarity of HVGs identified within each method, based on the variance-stabilising transformation method. **L)** Comparison of the average number of clusters detected in three comparable resolutions in each sample.

An inter-method comparison of the top 3,000 highly variable genes (HVGs) identified in each sample revealed that scRNA-seq samples consistently exhibit higher average Jaccard similarity scores (Figure 2K), suggesting that it is less affected by technical variability (such as site) than snRNA-seq (Supplementary Figure S6). Furthermore, across three comparable clustering resolutions applied to each sample, scRNA-seq consistently yields a higher number of clusters compared to snRNA-seq (Figure 2L), indicating its enhanced sensitivity to subtle transcriptional differences and ability to detect finer-scale cellular heterogeneity.

### Multi-sample Analyses

After per-sample QC (Supplementary Figure S8-S10), we merged the filtered cells from all samples to facilitate comprehensive downstream analyses between samples. Data were merged from a total of 80,422 nuclei and 91,633 whole cells, across 11 pairs of samples, into a single object comprising a total of 172,055 cells and 36,000 features. Following the removal of sample-specific genes, and normalisation, scaling and dimensionality reduction using the Seurat pipeline, UMAP visualisation revealed two distinct clusters corresponding to scRNA-seq and snRNA-seq cells respectively (Figure 3A-B). Having data generated across different laboratories and by different individuals allowed us to compare cells based on site, donor and method, and more accurately assess whether the observed differences between methods are systematic across sites, or instead the result of mishandling, in-house protocols, or instrument variability (Supplementary Figure S11).

**Figure 3.**
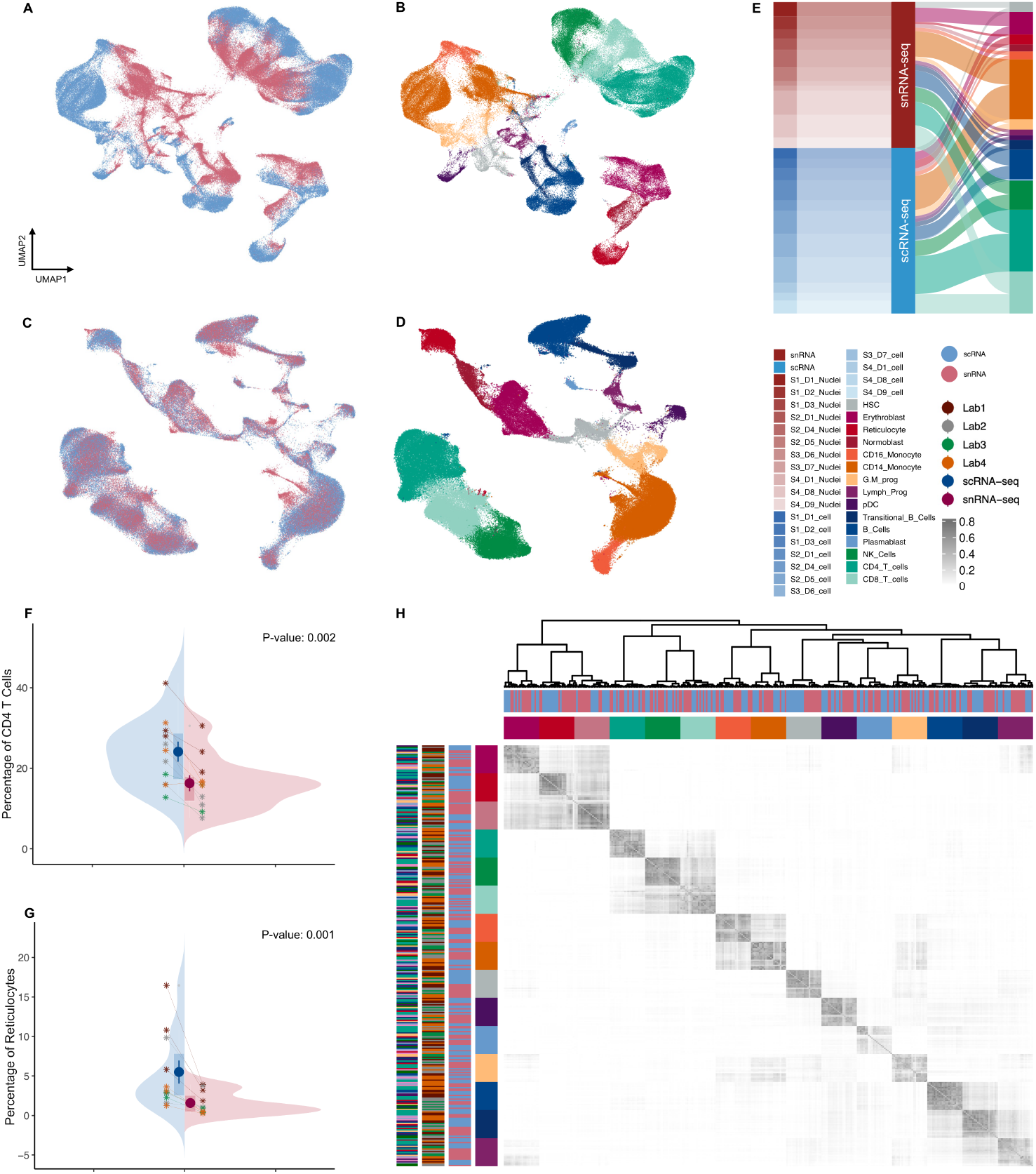
Integrated analysis of scRNA-seq and snRNA-seq datasets. **A–B)** UMAP visualisations of the merged datasets before batch correction, coloured by method (A) and cell type (B). **C–D)** UMAP visualisations of Harmony corrected embeddings, coloured by method (C) and cell type (D). **E)** Cell type composition across all 22 samples processed using the two approaches. **F–G)** Example split violin plots showing the proportion of CD4^+^ T cells (F) and Reticulocytes (G) within each matched sample pair. **H)** Heatmap and clustering of Jaccard similarity coefficients between marker genes for each cell type across individual samples.

Notably, this method-dependent separation appears to be primarily driven by differences in library complexity and the number of genes between the scRNA-seq and snRNA-seq data (Supplementary Figure S6). As illustrated in the UMAP visualisations (Figure 3C-D), correcting PCs for sample-specific effects using Harmony allowed integration of the data while preserving the biological variation among different cell types. Using these corrected PCs for clustering, we identified and annotated 15 distinct cell types. In spite of the inherent technical differences between scRNA-seq and snRNAseq, major cell populations are preserved across both sequencing approaches (Figure 3E). Subsequent pairwise comparisons of cell type proportions in each sample between the two methodologies revealed no significant differences in the composition of most cell types (Supplementary Figure S12), with the exception of erythroblasts, reticulocytes and CD4^+^ T cells (Figure 3F-G). Hierarchical clustering and visualization of similarity scores for the top 200 cell type–specific marker genes across 22 samples revealed that these markers clustered primarily according to the biological similarity of cell types and lineage proximity, rather than the capture methodology (Figure 3H), suggesting that the expression patterns of these markers are robust across both protocols, further underscoring the capability of each approach to detect all major cell types.

To investigate transcriptomic differences between scRNA-seq and snRNA-seq methodologies while accounting for cellular composition in each sample, we aggregated raw counts from the same cell types across the 11 samples per method. Pseudo-bulk differential expression analysis (DEA) (Table S1), showed that the number of up and down-regulated differentially expressed genes varied across cell types. While the numbers of upregulated and downregulated genes were comparable within each cell type, the average log-foldchange was consistently higher in genes up-regulated in scRNA-seq, suggesting that this method captures a greater dynamic range of gene expression (Figure 4A,B). To assess the similarity of these differentially expressed genes (DEGs) across cell types, we computed Jaccard similarity coefficients between the upregulated gene sets for each method. Hierarchical clustering of these coefficients demonstrated that cell types with close lineage proximity clustered together and exhibited higher overlap of DEGs. This grouping underscores the importance of accounting for cellular composition in single-cell data analyses. Interestingly, genes upregulated in scRNA-seq showed greater similarity across different cell types, mirroring a higher consistency previously observed in comparisons of highly variable genes (Figure 4C). Visualisation of gene length distributions among these DEG sets revealed that genes upregulated in snRNA-seq were predominantly longer, whereas those upregulated in scRNA-seq tended to be shorter (Figure 4D). This gene length bias indicates that the observed differences between the two technologies are likely influenced by technical effects rather than genuine biological variation. Therefore, we concluded that direct comparisons between scRNA-seq and snRNAseq datasets are most likely confounded by this bias and may lead to misleading interpretations. For example, we cannot determine with confidence whether the observed overexpression of stress-related genes in scRNA-seq reflects true biological stress induced during single-cell preparation or is instead an artifact of gene length bias. To more accurately compare the biological signal captured by each method, we performed a new round of pseudo-bulk DEA between selected cell-type pairs within each method separately (Supplementary Figure S13, S14). Specifically, in the CD4^+^ versus CD8^+^ T cell comparison (Figure 4E–F), most DEGs detected in snRNA-seq overlapped with those from the corresponding scRNA-seq analysis (Figure 4G), a substantial number of DEGs were specific to scRNA-seq. To further assess the biological relevance of these DEGs, we performed Gene Set Enrichment Analysis (GSEA) using immunologic signature gene sets. Notably, the top four enriched gene sets (adjusted *p<* 0.05, ranked by |NES|) were consistently enriched in both platforms, highlighting shared transcriptional programs underlying CD4^+^ and CD8^+^ T cell identity (Figure 4H).

**Figure 4.**
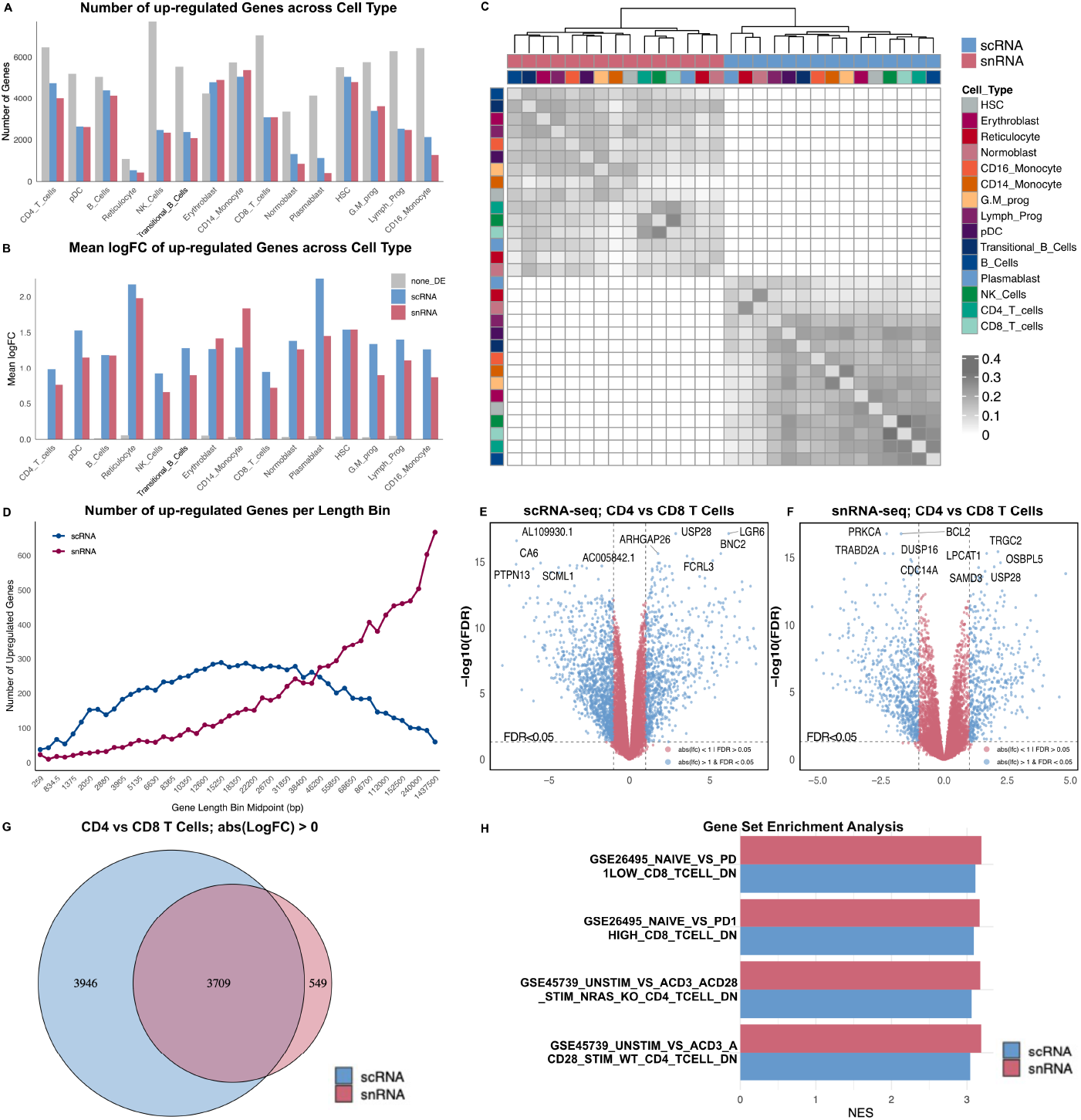
Differential gene expression (DE) analysis across scRNA-seq and snRNA-seq methodologies. **A–B)** Histograms showing results from pseudo-bulk DE analysis between scRNA-seq and snRNA-seq across 15 cell types. Grey bars indicate non-DE genes, blue bars genes upregulated in scRNA-seq, and red bars genes upregulated in snRNA-seq, grouped by cell type (A) and distribution of log fold-change values for each group (B). **C)** Jaccard similarity heatmap based on top 200 upregulated DEGs in each methodology, across different cell types. Samples are clustered using Ward.D2 on MDS-embedded Jaccard distances; annotations indicate cell type and methodology. **D)** Line plots show the number of upregulated genes in scRNA-seq and snRNA-seq datasets across midpoints of 50 gene-length bins. **E-F)** Volcano plots representing the distribution of DEGs and top marker genes for T cell comparisons within scRNA-seq (E) and snRNA-seq (F) **G)** Venn diagrams showing the overlap of DEGs between scRNA-seq and snRNA-seq in T cells. **H)** Gene Set Enrichment Analysis (GSEA) with the MSigDB C7 immunologic signature collection results. Bar plots show the normalised enrichment scores (NES) of the top 4 significantly enriched gene sets (adjusted *p<* 0.05, ranked by NES).

## Discussion

We observed lower sequencing saturation in scRNA-seq compared to snRNA-seq, combined with the overall higher number of gene counts and genes detected per cell, which suggests greater library complexity. This observation is consistent with findings from previous studies (Santiago et al., 2023; Mereu et al., 2020). Nuclei contain only a small fraction of the total cellular mRNA compared to the broader transcript pool captured by scRNA-seq, which includes both cytoplasmic and nuclear RNA. Although snRNA-seq typically yields lower read counts and detects fewer genes per nucleus, it has been shown to perform well in terms of capturing RNA molecules and sensitivity (Ding et al., 2020). The increased library complexity in scRNA-seq likely contributes to resolving more clusters, as noted in our results. As expected, we observed a higher proportion of reads mapping confidently to intronic regions in snRNA-seq data, in line with findings from previous studies (Truong et al., 2023; Habib et al., 2017; Kim et al., 2023). This high intronic content in nuclei may partially compensate for the loss of cytoplasmic RNA in snRNA-seq, and accounting for it may increase the number of detected clusters (Bakken et al., 2018). However, the higher intronic content in nuclear RNA may increase the likelihood of internal priming on premRNA, which could, in turn, contribute to a bias toward longer genes in snRNA-seq, as these genes are more likely to generate a greater number of Unique Molecular Identifiers (UMIs) (Gupta et al., 2022; Chamberlin et al., 2024). We observed a significant gene length bias between the two methodologies, with a bias toward the quantification of longer genes in snRNA-seq, consistent with findings from previous studies (Truong et al., 2023). In a study investigating gene length bias across single-cell technologies, Phipson *et al*. (Phipson et al., 2017) demonstrated that UMI-based singlecell protocols capture transcripts with relatively uniform gene lengths and are largely unaffected by gene length bias. Consistent with their findings, our results also showed minimal gene length bias in scRNA-seq data. However, upon removing a few highly expressed genes such as HBs and Malat1, we observed a noticeable bias toward longer genes in both scRNA-seq and snRNA-seq. This bias was more pronounced in the snRNA-seq data compared to scRNA-seq. Ambient RNA contamination poses a significant challenge, as it can lead to the false detection of highly abundant cytoplasmic genes in snRNA-seq across all cells, including those which are cell-type specific, thereby confounding downstream analyses (Young and Behjati, 2020). A higher percentage of ambient RNA observed in snRNA-seq could primarily be attributed to differences in sample preparation and the inherent variation of nuclear isolation processes. During snRNA-seq sample preparation, the lysis of cell membranes can release cytoplasmic RNA into the suspension, which may then be encapsulated within droplets, leading to contamination. Under-lysis may leave residual cytoplasmic RNA that contaminates nuclear preparations, while over-lysis can lead to RNA degradation and low recovery (Kim et al., 2023). We should also consider that the lower RNA content in nuclei, resulting in reduced library sizes, can exacerbate the percentage of ambient RNA contamination. Therefore, addressing ambient RNA contamination in snRNAseq requires a combination of optimised experimental protocols to minimise cytoplasmic RNA release, and using robust computational correction methods to ensure accurate and reliable transcriptomic profiling (Caglayan et al., 2022; Kersey et al., 2025). Our findings did not reveal a significant difference in doublet rates between the two methods. Notably, the doublet rate was correlated with the number of cells captured, aligning with the known relationship that higher cell loading densities can increase the probability of doublet occurrences (Germain et al., 2022).

The proportion of MT gene expression is a widely recognised QC metric in scRNA-seq analyses. Elevated MT gene percentages often indicate cellular stress or a broken membrane (Luecken and Theis, 2019). Overall, scRNA-seq data exhibited higher MT gene proportions than snRNA-seq due to the exclusion of cytoplasmic contents during the dissociation step in snRNA-seq (Kim et al., 2023; Truong et al., 2023). Despite the common practice of removing MT and RB genes from further analysis, and their frequent exclusion from quality control steps in snRNA-seq (Kim et al., 2023), our findings underscore the importance of assessing both the percentage and heterogeneity of these genes’ expression as key quality metrics. Such signals may highlight the more complex and delicate nature of the laboratory procedures involved in snRNA-seq data generation.

The UMAP visualisation ahead of removing batch effects, showed the cells obtained from scRNA-seq and snRNA-seq remained clearly segregated, aligning with other studies (Gupta et al., 2022; Truong et al., 2023). We observed that this distinction closely mirrors the pattern of library size difference on UMAPs, potentially downplaying concerns about the influence of other factors such as gene length bias. Merging of the cell atlases from both whole cells and nuclei resulted in well-integrated results in a previous benchmarking study (Mereu et al., 2020). Our results also showed that computational tools like Harmony are capable of integrating datasets obtained from these two methods (Korsunsky et al., 2019). However, due to the inherent under- or over-representation of certain cell types in each method, complementary analysis of each dataset may be more appropriate for addressing potential biases rather than integrative analysis (Kim et al., 2023). Differences in cell type compositions could be due to variations in cell viability during dissociation and cell shape and size affecting capture during the microfluidic step (Van Melkebeke et al., 2024). In our study, despite these differences as well as observed biases in library size and number of detected genes, both methods successfully captured all major cell types with only minor differences in composition of a few cell types. Notably, reticulocytes and erythroblasts were underrepresented in snRNA-seq and scRNA-seq datasets, respectively. Reticulocytes, which are enucleated or nearly so due to their maturation process into erythrocytes, lack nuclei and thus are less likely to be captured by snRNA-seq, which targets nuclear RNA. Conversely, erythroblasts, characterised by fragile membranes, high nuclear transcriptional activity, and abundant cytoplasmic HB transcripts (Schippel and Sharma, 2023; Liu et al., 2010) were less abundant in the scRNA-seq data. In particular, erythroblasts exhibited a distinct correlation between library size and the number of detected features compared to other cell types in the scRNA-seq data. In a study of two tissue types, neuroblastoma and metastatic breast cancer, immune cells were significantly more abundant in scRNA-seq across both tissue types, whereas parenchymal cells—particularly malignant cells—were more frequently captured by snRNA-seq (Slyper et al., 2020). Similarly, in a comparative analysis of matched normal and adenocarcinoma human lung samples, scRNA-seq detected a higher proportion of immune cells (81.5%), while snRNA-seq captured a greater proportion of epithelial cells (69.9%) (Renaut et al., 2024).

Previous studies comparing scRNA-seq and snRNA-seq have identified marker genes and gene sets that appear to be specific to each of these methodologies (Santo et al., 2025). For example, two independent reports have highlighted the overexpression of stress-related genes in scRNA-seq data (Slyper et al., 2020; Van Melkebeke et al., 2024). Our preliminary results from a pseudo-bulk differential expression analysis comparing transcriptomic profiles between the two methods revealed a pronounced gene length bias, with genes upregulated in snRNA-seq tending to be significantly longer. This observation aligns with recent findings by Chamberlin *et al*. (Chamberlin et al., 2024), and suggests that gene length is a major source of variation in cross-method comparisons. Addressing this challenge requires optimisation of experimental protocols in parallel with the development of normalisation methods that explicitly account for gene length. One such approach, based on normalising expression by the number of polyA regions per gene, has been proposed and applied in a recent study (Gupta et al., 2022).

While we have relatively well-developed lab protocols for both snRNA-seq and scRNA-seq, they are far from perfect. Some of the variation observed in our analysis is not due to the inefficacy of the sequencing technologies themselves, but rather it could be because of biases in the lab protocols for tissue processing, cell disaggregation, and preservation (Truong et al., 2023). For example, in snRNA-seq data, the per-cell proportions of MT and HB gene counts exhibited notable correlations across different sites. This emphasises the necessity of developing more efficient and reproducible protocols for laboratory procedures (Lee et al., 2023a). While there is no perfect method, using both techniques side by side can provide a more accurate view of the cell transcriptome (Santiago et al., 2023; Andrews et al., 2022). Pairing scRNA-seq with the measurement of other molecular layers like the epigenome, proteome, or spatial data could be another option to provide an integrative view on cellular identity (Deleersnijder et al., 2021; Hwang et al., 2022). Despite these inherent differences in laboratory workflows and transcript composition, when it comes to analysis, similar bioinformatics pipelines are typically used for the analysis of scRNAseq and snRNA-seq data (Kim et al., 2023). While selecting the appropriate assay is essential for generating high-quality results, the methods used for data processing play a critical role in shaping the final biological insights (Chamberlin et al., 2024). We recommend that bioinformaticians carefully consider the unique characteristics of data during analysis. In particular, the inherently lower library size and lower number of detected features in snRNA-seq should be taken into consideration when defining quality control thresholds. Background RNA contamination is also more pronounced in snRNA-seq and must be addressed during preprocessing. Additionally, the lower library complexity often results in fewer numbers of clusters at a given resolution compared to scRNA-seq. While MT transcripts are not typically expected in snRNA-seq data, their presence can still serve as a valuable quality indicator, potentially reflecting issues in the nuclei isolation procedure.

## Conclusion

In this study, we did not assume complete agreement between the results, nor was our aim to determine which method is superior. Instead, we conducted a series of analyses and visualisations to compare the outputs from two widely used single-cell RNA sequencing approaches on BMMCs. Although our batch correction and cell type annotations demonstrate that integrating datasets from these methods is technically feasible, we do not recommend directly comparing gene expression between the two methodologies. This caution is based on observed methodological biases—such as differences in library size, number of detected genes, and gene length—that may obscure biologically meaningful signals. Furthermore, this study offers practical guidance to researchers specifically bioinformaticians by highlighting key considerations related to the inherent characteristics of data generated by each methodology, helping inform decisions during preprocessing and downstream analysis. Finally, while both methods provide valuable information and have their own advantages and trade-offs, the choice between them should be guided by a well-defined study design that considers factors such as sample preservation method (fresh versus frozen), logistical constraints (e.g., timing), and study objectives (e.g., multimodal data).

## Methods

### Dataset

For this study, we re-used a publicly available dataset of bone marrow mononuclear cells (BMMCs), originally generated for a multi-modal single-cell benchmarking study (Figure 1A). In this dataset, biological samples from multiple healthy donors (aged 22–40) were processed across four different laboratories, where each sample was split into two parts, with one part subjected to transcriptomic profiling using 10x Multiome and the other using 10x CITE-seq. For both scRNAseq and snRNA-seq, 3’ gene expression libraries were sequenced on an Illumina NextSeq 2000 platform using paired-end reads with the following parameters: Read 1 (28 cycles), Index 1 (10 cycles), Index 2 (10 cycles), and Read 2 (90 cycles). The targeted sequencing depth was approximately 20,000 reads per cell or nucleus (Luecken et al., 2021). This dataset is among the largest and most comprehensive single-cell multi-modal benchmarking resources currently available, and we selected this based on several key considerations. First, public accessibility was a critical factor, ensuring that the data is freely available to the broader research community. Second, we prioritised a well-controlled experimental design to minimise batch effects. Third, the choice of tissue was critical. Bone marrow is a highly heterogeneous tissue containing diverse cell types at various stages of development. Its central role in hematopoiesis and immune system function makes it especially valuable for research in blood biology, immunology, hematologic diseases, and cancer. Additionally, extensive prior research and well-annotated reference datasets further support the utility of this tissue for benchmarking studies. Moreover, both datasets used in this study are multi-modal. The inclusion of a second modality introduces additional experimental challenges for each method but also enables cross-validation and provides an extra layer of information. The BAM files from 11 pairs of matched snRNA-seq and scRNA-seq samples were downloaded from NCBI under BioProject number PRJNA799242. These files were processed using various state-of-the-art bioinformatics tools, as illustrated in Figure 1B. Independent in-house data, used to further assess gene-length bias, are available in the Gene Expression Omnibus under accession number GSE303762.

### Sample-specific Analyses

#### Generating Count Matrices

Initially, the BAM files were converted to FASTQ format using the `bamtofastq` function from `Cell Ranger` version 7.1.0. The resulting FASTQ files were then demultiplexed and aligned to the GRCh38-2020-A-2.0.0 reference genome using the `cellranger -count` function with the `“include-introns”` option set to `true` to include intronic reads (10X Genomics, c). The generated count matrices served as input for downstream analyses conducted with the Seurat R package(Hao et al., 2021, 2024). We visualised and statistically compared the components of the Cell Ranger metrics summary report (10X Genomics, b) for each sample as demonstrated in Figure 2A.

#### Gene Annotation

Gene identifiers, coordinates and different gene-level attributes were retrieved using the `biomaRt` (R package version 2.58.2). Genomic length was defined as the distance between the start and end positions of each gene.

#### Ambient RNA

During sample preparation, broken and lysed cells release RNA into the surrounding solution. This extracellular background RNA, known as ambient RNA, can be encapsulated within droplets and subsequently sequenced alongside cellular transcripts, introducing background noise in downstream analyses. The `SoupX` (version 1.6.2) R package was used to estimate and remove the ambient RNA from each individual sample (Young and Behjati, 2020). To provide clustering labels for the `setClusters` function, each individual dataset was normalised, and variable feature selection and scaling were performed using Seurat’s `SCTransform` function. Principal components (PCs) were calculated from the top 3000 variable features. Seurat’s `FindNeighbors` and `FindClusters` functions were used to build the graph and cluster cells based on the Louvain algorithm, respectively.

#### Doublet Detection

Doublets occur when two individual cells or nuclei are inadvertently captured within the same droplet, resulting in combined transcripts from both cells that can confound downstream analyses. The `scDblFinder` R package (version 1.16.0) (Germain et al., 2022) was used to detect doublets. To improve detection accuracy, after removing ambient RNA, a second round of normalisation and clustering was performed, and the resulting cluster labels were then supplied to scDblFinder.

#### Biology Aware QC

Upon observing significant differences in library size and feature counts between scRNA-seq and snRNA-seq methods, combined with the high heterogeneity of bone marrow mononuclear cells (BMMCs), we anticipated natural variability in quality control (QC) metrics across samples. To ensure a fair and biologically meaningful comparison, we employed an adaptive, data-driven approach to identify low-quality cells at the cluster or cell-type level. This strategy accounts for biological variability and has been demonstrated to preserve rare cell populations with low library sizes (Subramanian et al., 2022). After a third round of clustering, we identified and removed outlier cells within each cluster whose QC metrics deviated beyond ±3 median absolute deviations (MAD) from the cluster median for library size and number of features per cell. Finally, we applied shallow, uniform thresholds across all samples to exclude cells with extreme QC metrics: `nCount_RNA` *<* 100, `nFeature_RNA` *<* 50, `percent.mt` *>* 10, and removal of detected doublets.

#### Highly Variable Gene Consistency

Feature selection as a first step in dimensionality reduction has an important role in clustering results (Luecken and Theis, 2019). To evaluate the consistency and method-specific differences in HVGs composition that underlie downstream dimensionality reduction and clustering, we compared the top 3,000 HVGs identified using two different approaches: SCTransform v2 and Seurat’s variance stabilising transformation (VST) method. We assessed the overlap between these HVG sets using the Jaccard similarity index, providing a quantitative measure of similarity.

### Multi-sample Analyses

After merging the individual Seurat objects, to reduce potential confounding effects, we excluded specific gene sets known to introduce bias or noise. Biological sex genes located on chromosomes X and Y were excluded alongside other known confounding genes, including immunoglobulin, T-cell receptor, HB, RB, MT, and MALAT1. Then we subjected the merged dataset to a standard preprocessing workflow using Seurat’s default parameters. This included normalisation of the raw count data with the `NormalizeData()` function, identification of 3000 HVGs via `FindVariableFeatures()`, and scaling of the data using `ScaleData()`. Considering the differentiating processes in BMMCs, we did not want to regress out all cell cycle signals, as this could make it challenging to distinguish between hematopoietic stem cells as non-cycling (quiescent) cells and progenitors as cycling (dividing) cells. Therefore, to preserve the signals separating non-cycling cells from cycling cells, we only regress out the differences between the G2M and S phase scores, as computed by the `CellCycleScoring()` function in Seurat, as recommended in Seurat Cell-Cycle Scoring Vignette (Satija et al., 2021; Hao et al., 2021). Subsequently, we performed dimensionality reduction through principal component analysis (PCA) using the `RunPCA()` function.

#### Batch Effect Removal

To correct PCA embeddings for batch effects, we employed the `Harmony` R package (version 1.2.0), where 22 sample ids were provided as the variable to remove (Korsunsky et al., 2019). A shared nearest neighbour (SNN) graph was constructed using Seurat’s `FindNeighbors()` function, and clustering was performed using the Louvain algorithm via the `FindClusters()` function across a range of resolutions. Uniform Manifold Approximation and Projection (UMAP) embeddings were computed using the `RunUMAP()` function for both Harmony-corrected and uncorrected PCs to facilitate visualisation.

#### Consensus Cell Type Annotation

We employed a consensus-based cell type annotation strategy that integrates multiple complementary data sources. Initially, we utilised the clustree R package (version 0.5.1) (Zappia and Oshlack, 2018) to visualise hierarchical relationships and potential lineage trajectories among clusters across varying resolutions, aiding in the identification of resolution with stable clustering. To examine inter-cluster relationships and overall data structure, we visualised UMAP embeddings. Subsequently, we assessed the expression patterns of canonical cell type-specific marker genes using Seurat’s `DotPlot()` function (Stuart et al., 2019). Additionally, annotations from the original studies of public datasets were incorporated to validate our cell type assignments(Luecken et al., 2021). We visualised the proportion of these independent cell labels within each cluster using heatmaps. Finally, our annotations were confirmed and refined by an expert as a gold standard (Clarke et al., 2021).

#### Cell Type–Specific Marker Gene Consistency

To evaluate the consistency of cell–type–specific marker genes across samples and between the two methodologies, we performed differential expression analysis independently for each sample using Seurat’s `FindAllMarkers()` function. For each cell type within a sample, we identified marker genes by comparing the expression profiles of that cell type against all other cells in the same sample. To assess the similarity of marker genes across samples and methods, we computed pairwise Jaccard similarity scores based on the top 200 marker genes (adjusted *p*-value *<* 0.05 and log_2_ fold change *>* 1) for each cell type within each sample.

#### Hierarchical Clustering

The similarity matrices were visualised using the `pheatmap` R package (version 1.0.12). To facilitate hierarchical clustering, each similarity matrix was transformed into a distance matrix using the formula: `distance` = 1 − `similarity`. Classical multidimensional scaling (MDS) was then applied to embed the distance matrix into Euclidean space. Hierarchical clustering was performed on the MDS coordinates using the `Ward.D2` linkage method, as implemented in the `pheatmap` package.

#### Pseudo-bulk Differential Expression Analysis

To more accurately identify genes differentially expressed between two approaches while accounting for heterogeneity in cell type compositions across samples, we performed pseudo-bulk differential expression analysis at the cell type level. For each of the 22 samples, raw counts from cells of the same type were aggregated to create pseudo-bulk profiles. Differential expression analysis was then conducted using the `edgeR` R package (version 4.0.16). A design matrix was constructed to model both the experimental methodology and the paired nature of the samples. In total, we carried out 15 differential expression analyses corresponding to the distinct cell types identified. To evaluate the overlap of DEGs across these comparisons, we calculated the Jaccard similarity coefficient and the results were visualised using heatmaps. To mitigate the effect of gene length bias, we performed an additional round of pseudo-bulk differential expression analysis to identify genes that distinguish CD4 and CD8 T cells within each approach. To evaluate the biological relevance of these DEGs, we conducted Gene Set Enrichment Analysis (GSEA) using the `clusterProfiler` R package (version 4.10.1). Genes from the CD4 vs CD8 T cell comparisons were ranked by log fold change, and GSEA was performed separately for the scRNA-seq and snRNA-seq datasets using the C7 immunologic signature gene sets from the Molecular Signatures Database (MSigDB), accessed via the `msigdbr` R package (version 7.5.1). Significantly enriched gene sets were defined as those with an adjusted *p*-value *<* 0.05.

#### Statistical Testing

A paired Wilcoxon signed-rank test was conducted using R’s `wilcox.test()` function from the base `stats` package to assess differences between sequencing methods within matched samples. The analysis was performed on various gene attribute categories and the relative proportions of cell types across these paired samples.

The online version contains supplementary plots and tables available at https://github.com/GhamsariReza/snRNA_vs_scRNA_comparison.

## Supporting information

Extended Format

## Abbreviations

BMMCs: Bone Marrow Mononuclear Cells
snRNA-seq: Single Nucleus RNA Sequencing
scRNA-seq: Single Cell RNA Sequencing
DEGs: Differentially Expressed Genes
DEA: Differential Expression Analysis
GSEA: Gene Set Enrichment Analysis
HVGs: Highly Variable Genes
MAD: median absolute deviation
QC: Quality Control
MT: Mitochondrial
RB: Ribosomal
HBs: Hemoglobins
UMIs: Unique Molecular Identifiers
SNN: Shared Nearest Neighbour
UMAP: Uniform Manifold Approximation and Projection
PCA: Principal Component Analysis

## Declarations

### Ethics Approval

This study did not involve any experiments on human participants or animals. All analyses were performed using publicly available datasets, and ethics approval was not required.

### Consent for publication

All authors read and approved the final manuscript.

### Availability of Data and Materials

All code used to reproduce the results and figures in this study is available at: https://github.com/GhamsariReza/snRNA_vs_scRNA_comparison. The study was conducted using publicly available datasets, which can be accessed at: https://www.ncbi.nlm.nih.gov/bioproject/PRJNA799242.

## Authors’ Contributions

R.G. conceived the study idea, performed the data analysis, and wrote the manuscript. M.R. and H.K. provided supervision and guidance throughout the project and critically revised the manuscript. C.G., R.T. and Y.Y. contributed ideas and analysis input at various stages of the project. N.L. revised the manuscript.

## Acknowledgements

We gratefully acknowledge the Research Computing Team and IT Support at the Walter and Eliza Hall Institute for providing high-performance computing (HPC) resources and technical assistance. We also thank Dr Ashleigh Solano, Dr Davide Vespasiani, Dr Hamish King and Dr Nona Farbehi for their valuable feedback on this project.

