## Extended Format for "Comparative Analysis of Single-Nucleus and Single-Cell RNA Sequencing in Human Bone Marrow Mononuclear Cells: Methodological Insights and Trade-offs"

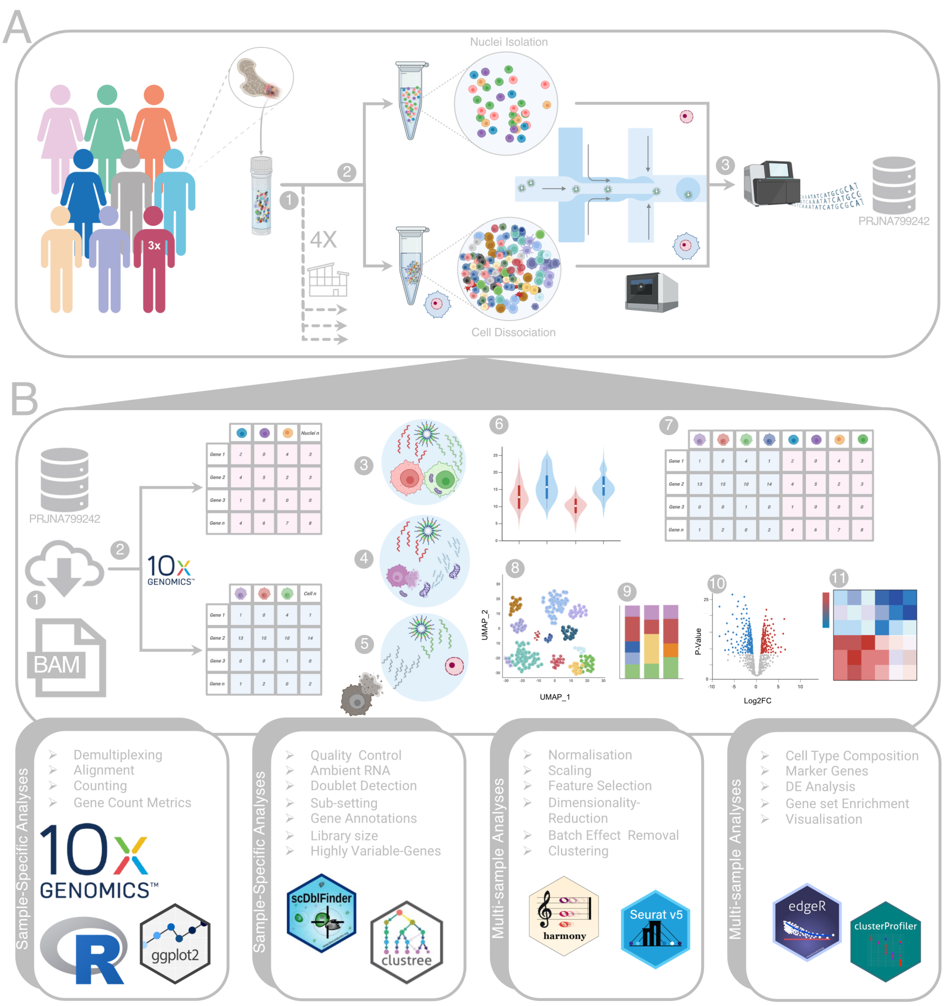


**Figure 1. Schematic representation of the study design, dataset and analysis workflow. A)** This study re-analysed a publicly available dataset comprising 11 matched pairs of BMMCs samples from healthy donors. These samples were processed across four (A1) different laboratories, where each donor's sample was divided into two aliquots: one processed using scRNA-seq and the other using snRNA-seq (A2). Libraries for both scRNA-seq and snRNA-seq were sequenced on the Illumina NextSeq 2000 platform (A3). **B)** Comparative analyses between the two approaches were conducted using the Seurat pipeline and other R packages. For sample-specific analyses, the raw data were downloaded (B1) and aligned to the reference genome using 10x Genomics' Cell Ranger software (B2). The QC steps include doublet detection (B3), removing cells with high mitochondrial gene percentage (B4), ambient RNA removal (B5) and a biology-aware QC approach (B6), which was performed first on each sample separately. Multi-sample analyses, followed by merging different samples (B7), applying batch effect removal, clustering (B8), cell type annotation (B9), differential expression (DE) analysis (B10) and subsequent downstream analyses (B11). Figure created with `BioRender.com`.

cell types and lineage proximity, rather than the capture methodology (Figure 3H), suggesting that the expression patterns of these markers are robust across both protocols, further underscoring the capability of each approach to detect all major cell types.

To investigate transcriptomic differences between scRNA-seq and snRNA-seq methodologies while accounting for cellular composition in each sample, we aggregated raw counts from the same cell types across the 11 samples per method. Pseudo-bulk differential expression analysis (DEA) (Table S1), showed that the number of up and down-regulated differentially expressed genes varied across cell types. While the numbers of upregulated and downregulated genes were comparable within each cell type, the average log-fold-change was consistently higher in genes up-regulated in scRNA-seq, suggesting that this method captures a greater dynamic range of gene expression (Figure 4A,B). To assess the similarity of these differentially expressed genes (DEGs) across cell types, we computed Jaccard similarity coefficients between the upregulated gene sets for each method. Hierarchical clustering of these coefficients demonstrated that cell types with close lineage proximity clustered together and exhibited higher overlap of DEGs. This grouping underscores the importance of accounting for cellular composition in single-cell data analyses. Interestingly, genes upregulated in scRNA-seq showed greater similarity across different cell types, mirroring a higher consistency previously observed in comparisons of highly variable genes (Figure 4C). Visualisation of gene length distributions among these DEG sets revealed that genes upregulated in snRNA-seq were predominantly longer, whereas those upregulated in scRNA-seq tended to be shorter (Figure 4D). This gene length bias indicates that the observed differences between the two technologies

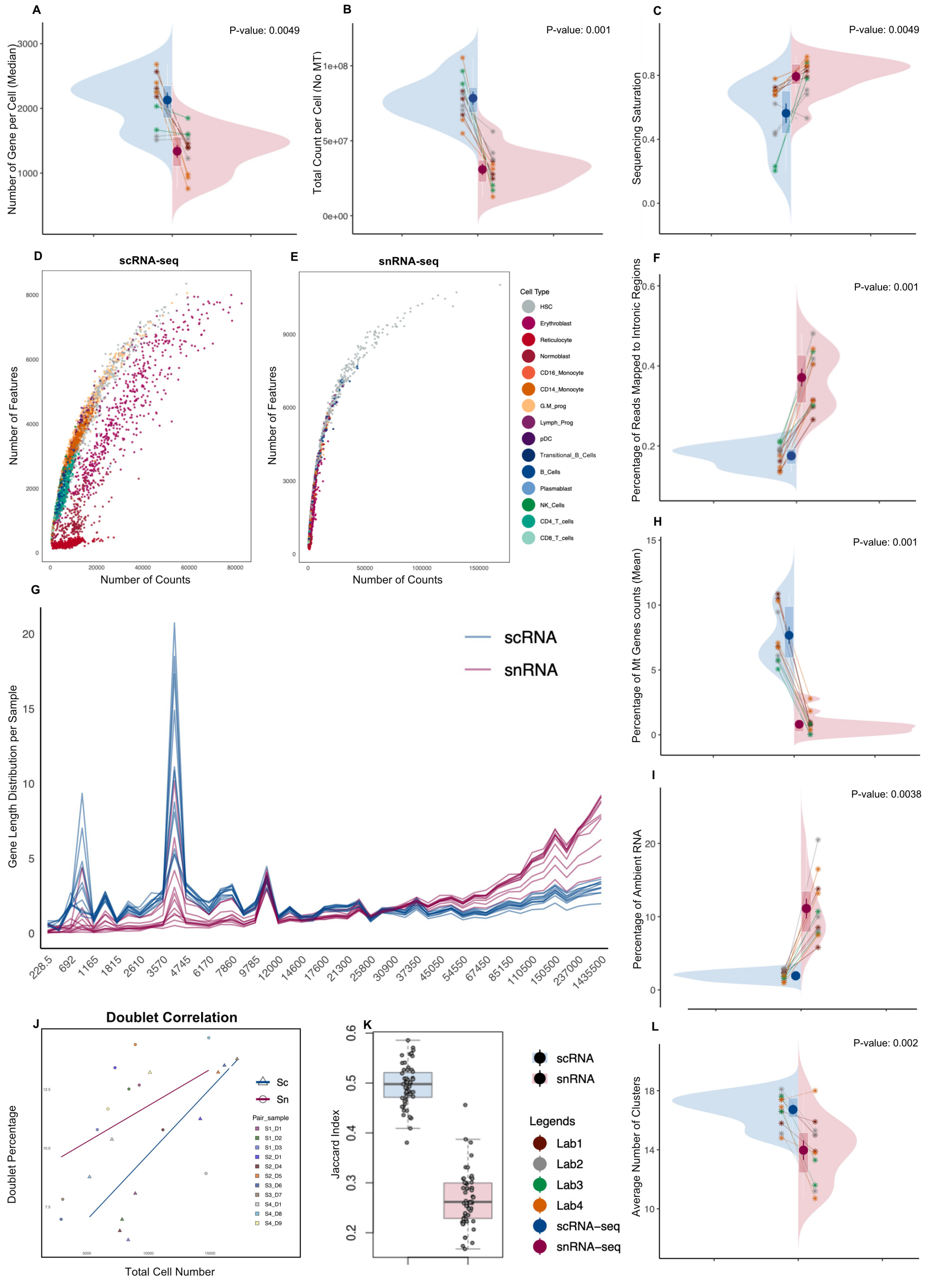

**Figure 2. Sample-level comparisons between scRNA-seq and snRNA-seq. A–C)** Split violin plots display the distribution of key sequencing and cell quality metrics, with red representing snRNA-seq and blue representing scRNA-seq. Each pair of samples is connected by a line. Samples are coloured by the laboratory to indicate batch origin (A, B, C, F, H, I, L). Median number of genes detected per cell for each sample (A), Library size per sample after removing MT genes (B) and Sequencing saturation (C). **D–E)** Scatter plots showing an example of the correlation between library size and number of genes per cell in scRNA-seq (D) and snRNA-seq (E), coloured by cell type. Scatter plots for all samples are shown in Figure S7. **F)** Split violin plots showing proportion of reads mapped to intronic regions per sample. **G)** Distribution of total UMI counts per gene length bin across 50 gene length bins across all 22 samples. **H–I)** Split violin plots comparing the percentage of MT gene content (H) and ambient RNA (I). **J)** Correlation between the percentage of detected doublets and the total number of cells recovered per sample. **K)** Boxplots showing Jaccard scores representing similarity of HVGs identified within each method, based on the variance-stabilising transformation method. **L)** Comparison of the average number of clusters detected in three comparable resolutions in each sample.

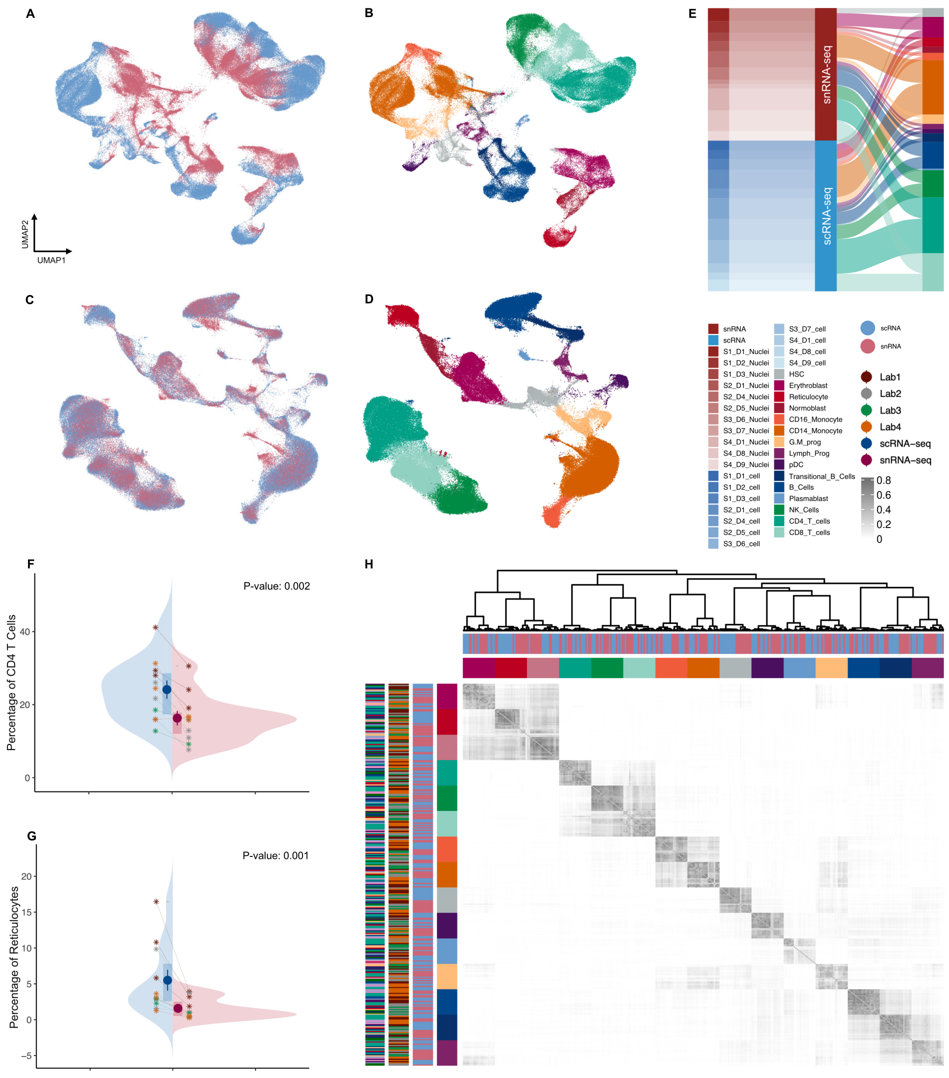


**Figure 3. Integrated analysis of scRNA-seq and snRNA-seq datasets. A–B)** UMAP visualisations of the merged datasets before batch correction, coloured by method (A) and cell type (B). **C–D)** UMAP visualisations of Harmony corrected embeddings, coloured by method (C) and cell type (D). **E)** Cell type composition across all 22 samples processed using the two approaches. **F–G)** Example split violin plots showing the proportion of CD4$^{+}$ T cells (F) and Reticulocytes (G) within each matched sample pair. **H)** Heatmap and clustering of Jaccard similarity coefficients between marker genes for each cell type across individual samples.

are likely influenced by technical effects rather than genuine biological variation. Therefore, we concluded that direct comparisons between scRNA-seq and snRNA-seq datasets are most likely confounded by this bias and may lead to misleading interpretations. For example, we cannot determine with confidence whether the observed overexpression of stress-related genes in scRNA-seq reflects true biological stress induced during single-cell preparation or is instead an artifact of gene length bias. To more accurately compare the biological signal captured by each method, we performed a new round of pseudo-bulk DEA between selected cell-type pairs within each method separately (Supplementary Figure S13, S14). Specifically, in the CD4$^{+}$ versus CD8$^{+}$ T cell comparison (Figure 4E–F), most DEGs detected in snRNA-seq overlapped with those from the corresponding scRNA-seq analysis (Figure 4G), a substantial number of DEGs were specific to scRNA-seq. To further assess the biological relevance of these DEGs, we performed Gene Set Enrichment Analysis (GSEA) using immunologic signature gene sets. Notably, the top four enriched gene sets (adjusted $p < 0.05$, ranked by |NES|) were consistently enriched in both platforms, highlighting shared transcriptional programs underlying CD4$^{+}$ and CD8$^{+}$ T cell identity (Figure 4H).

#### Discussion

We observed lower sequencing saturation in scRNA-seq compared to snRNA-seq, combined with the overall higher number of gene counts and genes detected per cell, which suggests greater library complexity. This observation is consistent with findings from previous studies (Santiago et al., 2023; Mereu et al., 2020). Nuclei contain only a small fraction of the total cellular mRNA compared to the broader transcript pool captured by scRNA-seq, which includes both cytoplasmic and nu-

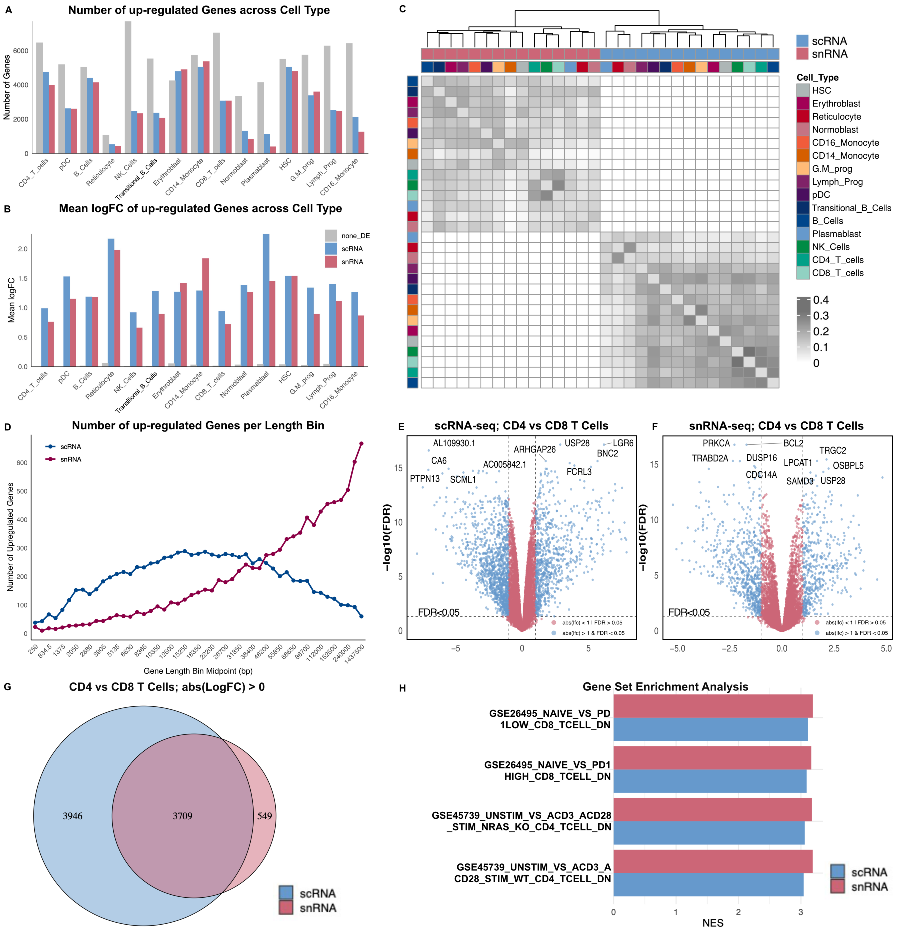


**Figure 4. Differential gene expression (DE) analysis across scRNA-seq and snRNA-seq methodologies. A–B)** Histograms showing results from pseudo-bulk DE analysis between scRNA-seq and snRNA-seq across 15 cell types. Grey bars indicate non-DE genes, blue bars genes upregulated in scRNA-seq, and red bars genes upregulated in snRNA-seq, grouped by cell type (A) and distribution of log fold-change values for each group (B). **C)** Jaccard similarity heatmap based on top 200 upregulated DEGs in each methodology, across different cell types. Samples are clustered using Ward.D2 on MDS-embedded Jaccard distances; annotations indicate cell type and methodology. **D)** Line plots show the number of upregulated genes in scRNA-seq and snRNA-seq datasets across midpoints of 50 gene-length bins. **E-F)** Volcano plots representing the distribution of DEGs and top marker genes for T cell comparisons within scRNA-seq (E) and snRNA-seq (F) **G)** Venn diagrams showing the overlap of DEGs between scRNA-seq and snRNA-seq in T cells. **H)** Gene Set Enrichment Analysis (GSEA) with the MSigDB C7 immunologic signature collection results. Bar plots show the normalised enrichment scores (NES) of the top 4 significantly enriched gene sets (adjusted $p < 0.05$, ranked by NES).

#### Competing interests

The authors declare no competing interests.

#### Funding

R.G. is supported by the Australian government Research Training Program (RTP) scholarship and partially funded by a PhD top-up scholarship from the Engineering Top-Up Award (ETA), School of Biomedical Engineering, UNSW, Sydney, Australia, as well as a Tour de Cure PhD grant. M.R. is supported by Australian National Health and Medical Research Council (NHMRC) Investigator Grant (GNT2017257), Victorian State Government Operational Infrastructure Support, Australian Government NHMRC IRIISS and support from the Australian Cancer Research Foundation.

#### Authors' Contributions

R.G. conceived the study idea, performed the data analysis, and wrote the manuscript. M.R. and H.K. provided supervision and guidance throughout the project and critically revised the manuscript. C.G., R.T. and Y.Y. contributed ideas and analysis input at various stages of the project. N.L. revised the manuscript.

#### Acknowledgements

We gratefully acknowledge the Research Computing Team and IT Support at the Walter and Eliza Hall Institute for providing high-performance computing (HPC) resources and technical assistance. We also thank Dr Ashleigh Solano, Dr Davide Vespasiani, Dr Hamish King and Dr Nona Farbehi for their valuable feedback on this project.

### Supplementary Information

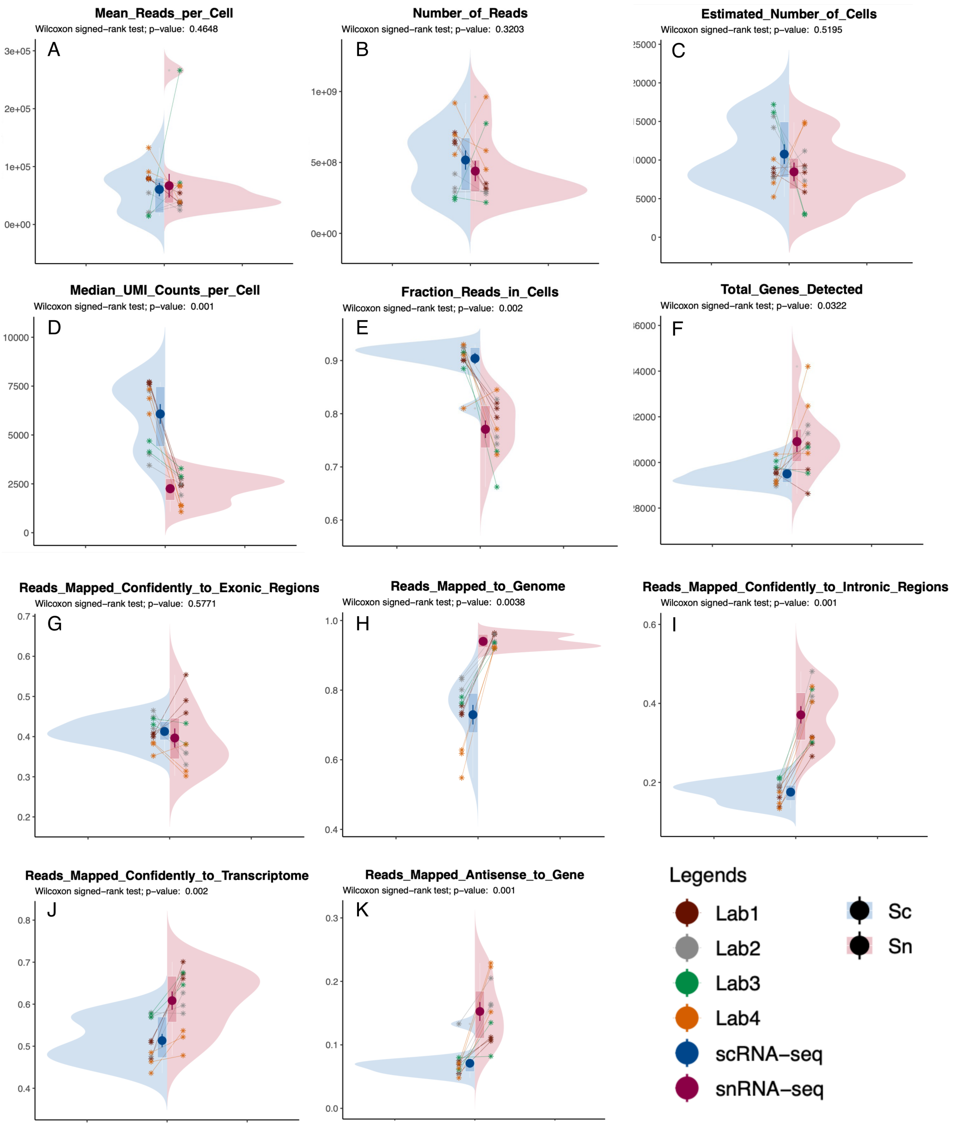


**Figure S1.** Split violin plots comparing various attributes from the Cell Ranger summary reports. Individual-level comparisons between scRNA-seq and snRNA-seq.

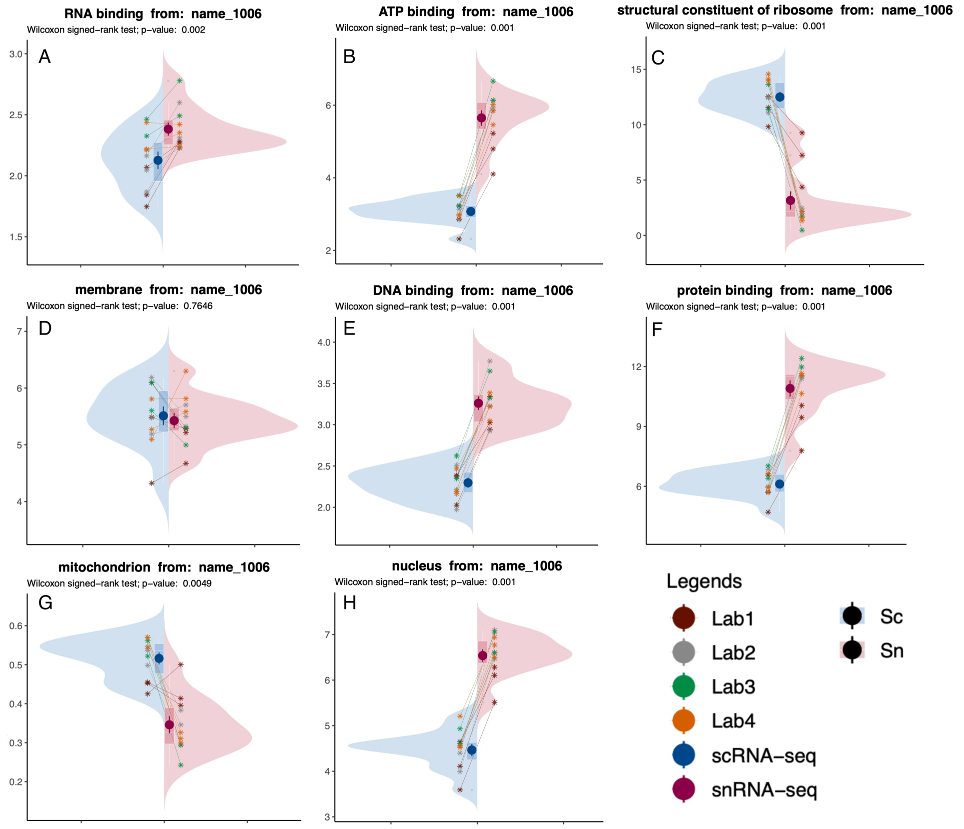


**Figure S2.** Split violin plots comparing various attributes associated with the GO term name_1006 retrieved using the `biomaRt` R package.

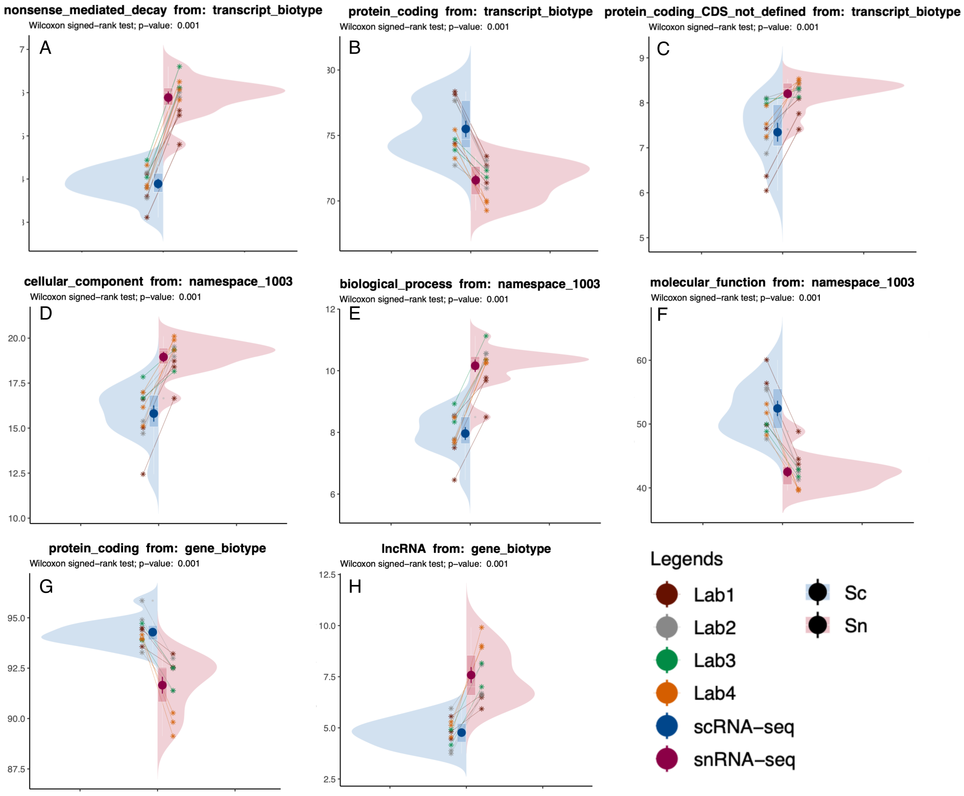


**Figure S3.** Split violin plots comparing various attributes associated with the GO terms. **A–C)** Transcript biotype; **D–F)** namespace_1003 **G–H)** Gene biotype. All attributes were retrieved using the `biomaRt` R package.

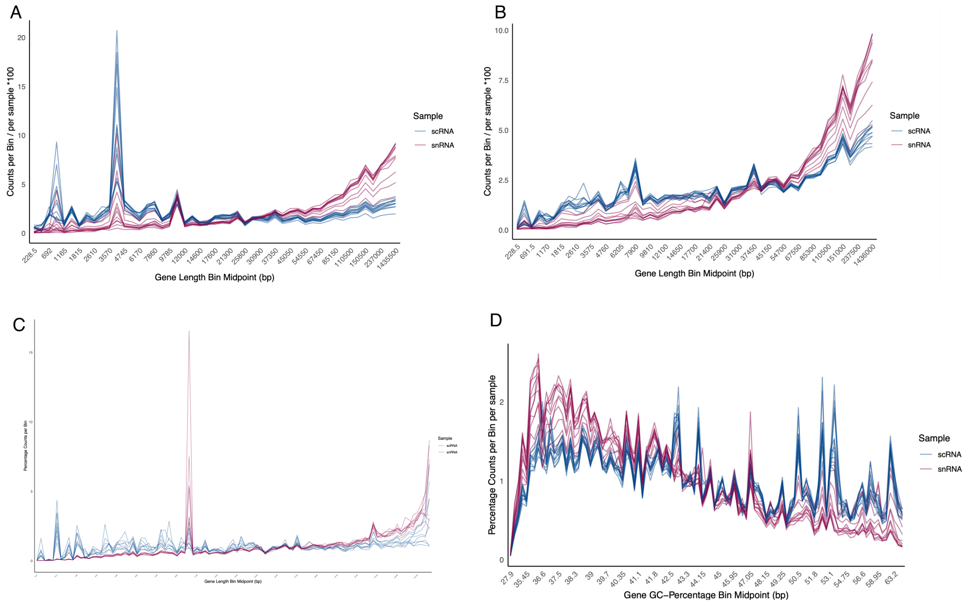


**Figure S4.** Distribution plots comparing gene length and GC content attributes retrieved using the `biomaRt` R package. **A)** Distribution of total UMI counts per gene length bin across 50 bins for all 22 samples, including all genes. **B)** Distribution of total UMI counts per gene length bin across 50 bins for all 22 samples, after removing HBs, RBs, *Malat1*, and MT genes. **C)** Distribution of total UMI counts per gene length bin across 50 bins for two additional independent in-house samples at the cluster level. Each sample contains 8 clusters representing different cell lines, pooled and sequenced using either snRNA-seq or scRNA-seq. **D)** Distribution of total UMI counts per gene GC content bin across 50 bins for all 22 samples, after removing HBs, RBs, *Malat1*, and MT genes.

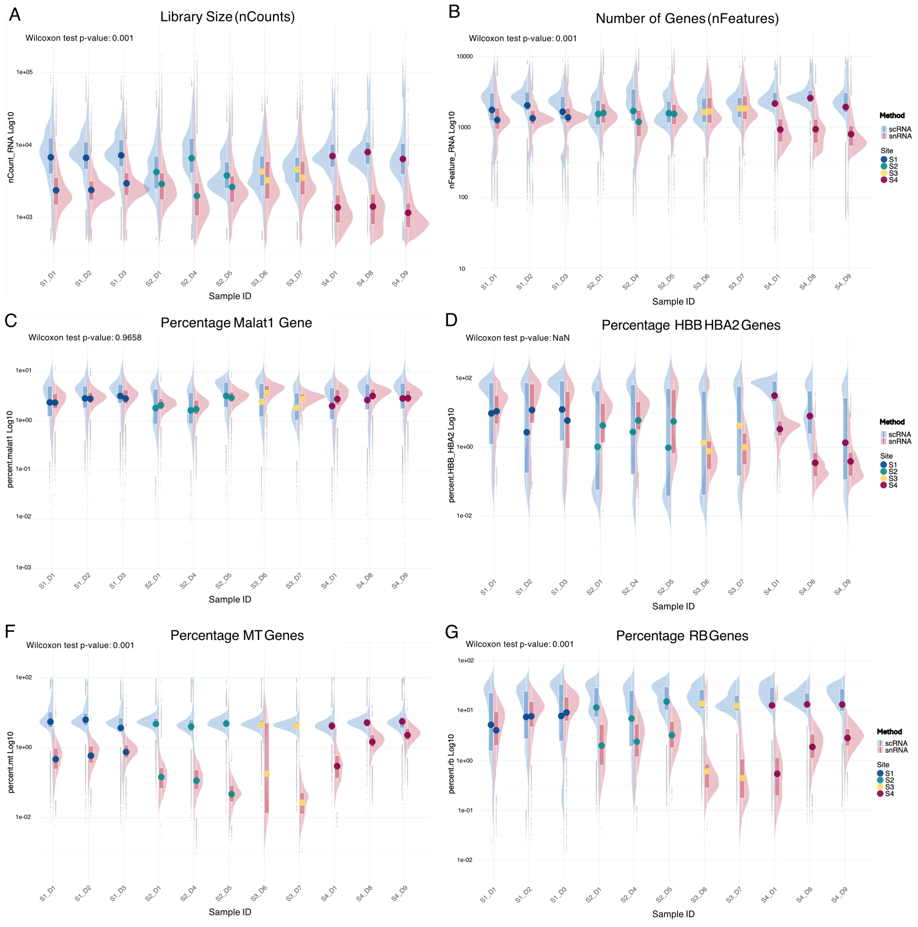


**Figure S5.** Split violin plots comparing various quality indices for individual cells across different paired samples.

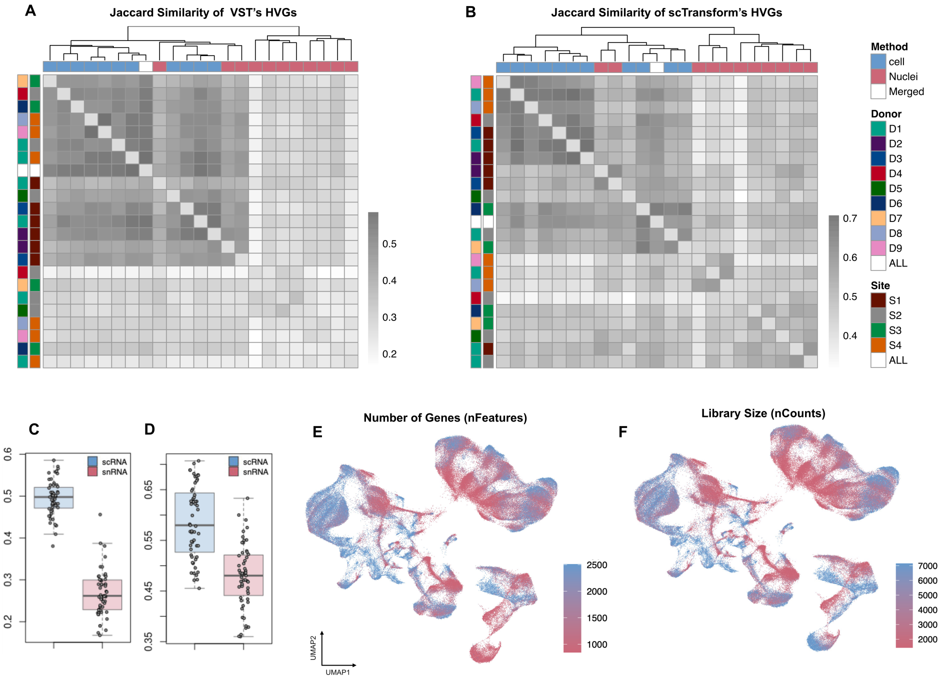


**Figure S6. A-B)**Heatmap and hierarchical clustering of Jaccard similarity coefficients between sets of highly variable genes identified using **(A)** Variance Stabilising Transform (VST) and **(B)** SCTransform, for 22 individual samples across the two methodologies, as well as the merged object ("Both"). **C–D)** Boxplots showing Jaccard similarity scores for highly variable genes (HVGs) within each method, calculated using variance stabilising transformation (C) and SCTransform (D). **E–F)** Feature plots showing the distribution of the number of genes (E) and library size (F) over the UMAP embedding.

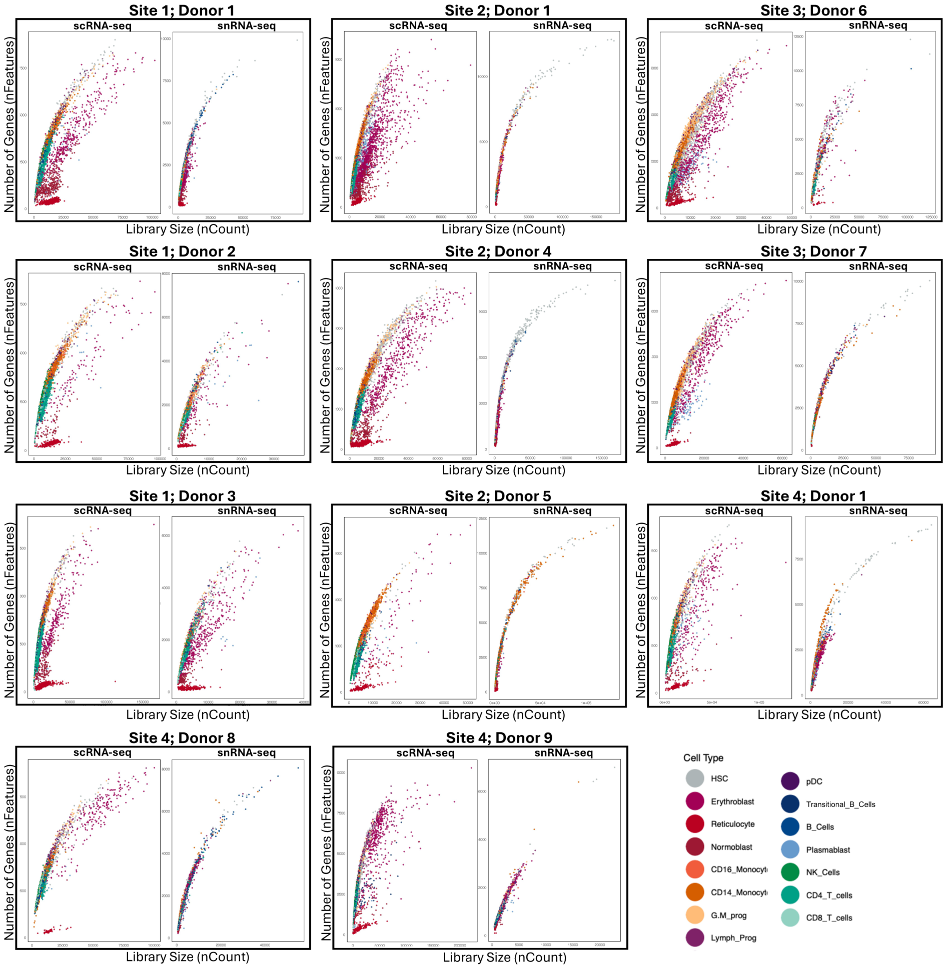


**Figure S7.** Scatter plots showing the correlation between library size and the number of detected genes per cell across eleven pairs of samples processed using scRNA-seq and snRNA-seq.

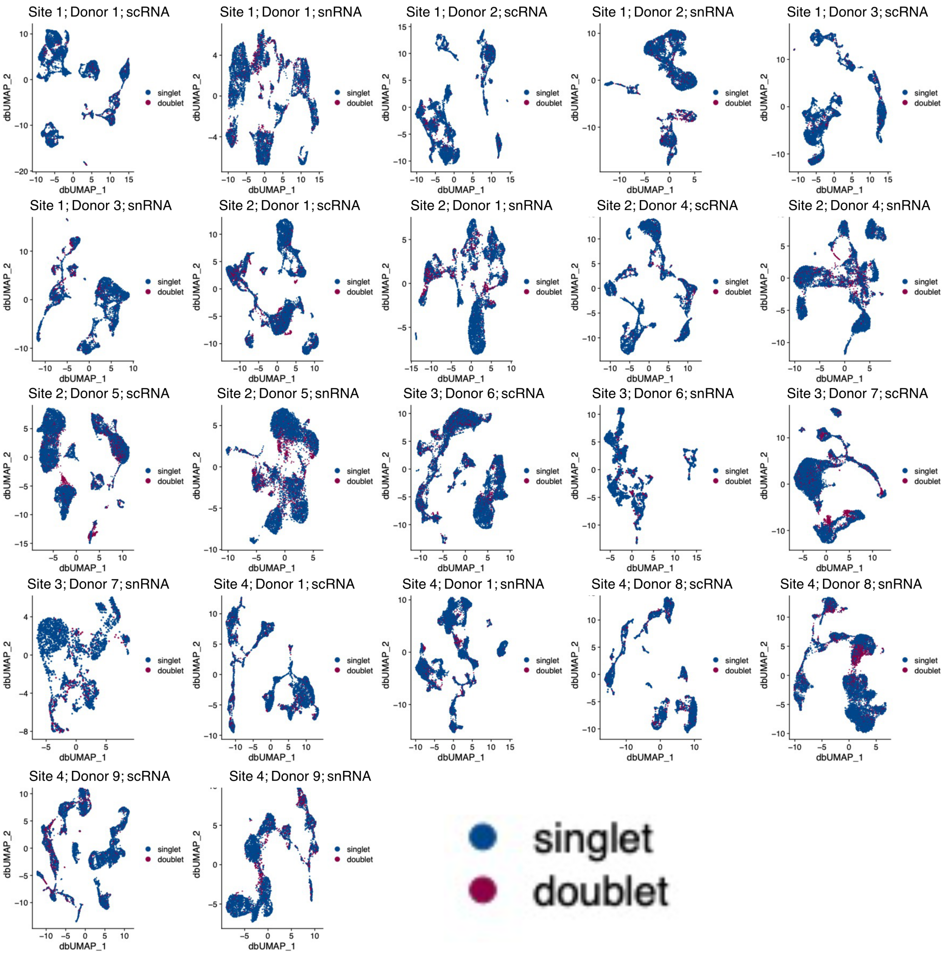


**Figure S8.** UMAP visualisation of detected doublets across 22 samples at the individual-level analysis.

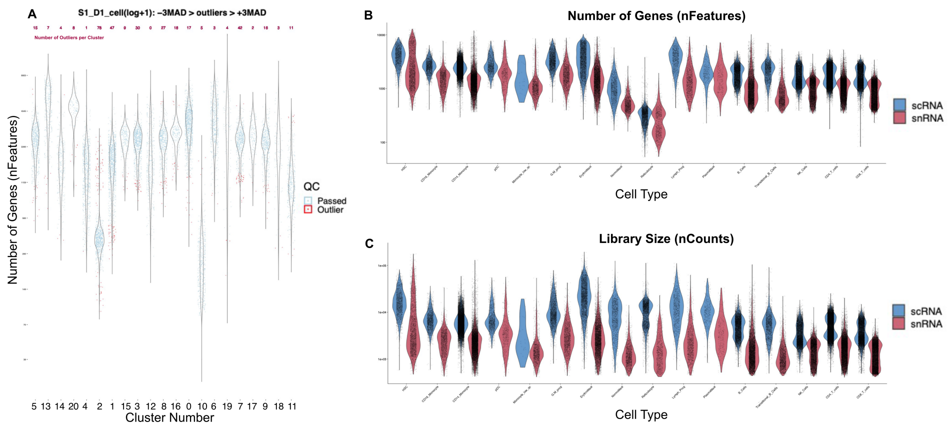


**Figure S9. A)** Example violin plot demonstrating biologically informed quality control in a single sample, where outliers are identified using a $\pm 3$ MAD threshold within each cluster. A PDF file containing corresponding plots for all samples and features is available for download on the GitHub page. **B–C)** Paired violin plots showing: **(B)** Number of detected features (genes) per cell across different cell types in the merged dataset, split and coloured by sequencing method (scRNA-seq vs. snRNA-seq). **(C)** Library size (total counts per cell) across different cell types in the merged dataset, also split and coloured by sequencing method.

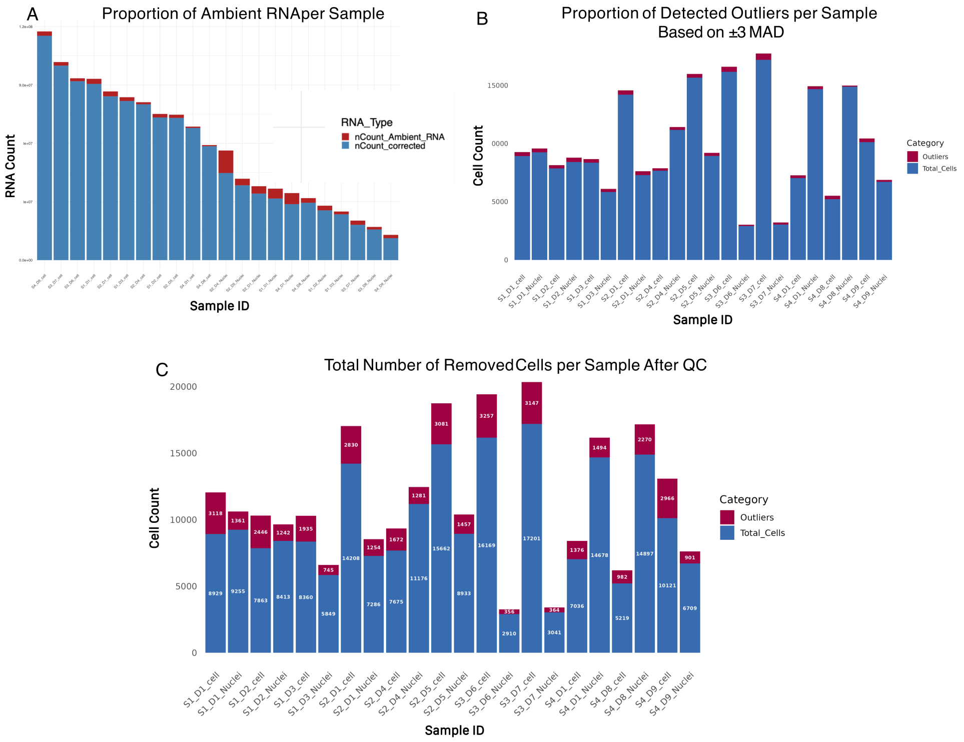


**Figure S10.** Stacked bar plots representing: **A)** Proportion of ambient RNA in each sample. **B)** Proportion of cells identified as outliers based on a $\pm 3$ MAD threshold within each cluster. **C)** Proportion of low-quality cells that passed the following quality filters: `nCount_RNA > 100 && nFeature_RNA > 50 && percent.mt < 10 && scDblFinder_RNA_SCT == "singlet" && QC_MAD == "Passed"`.

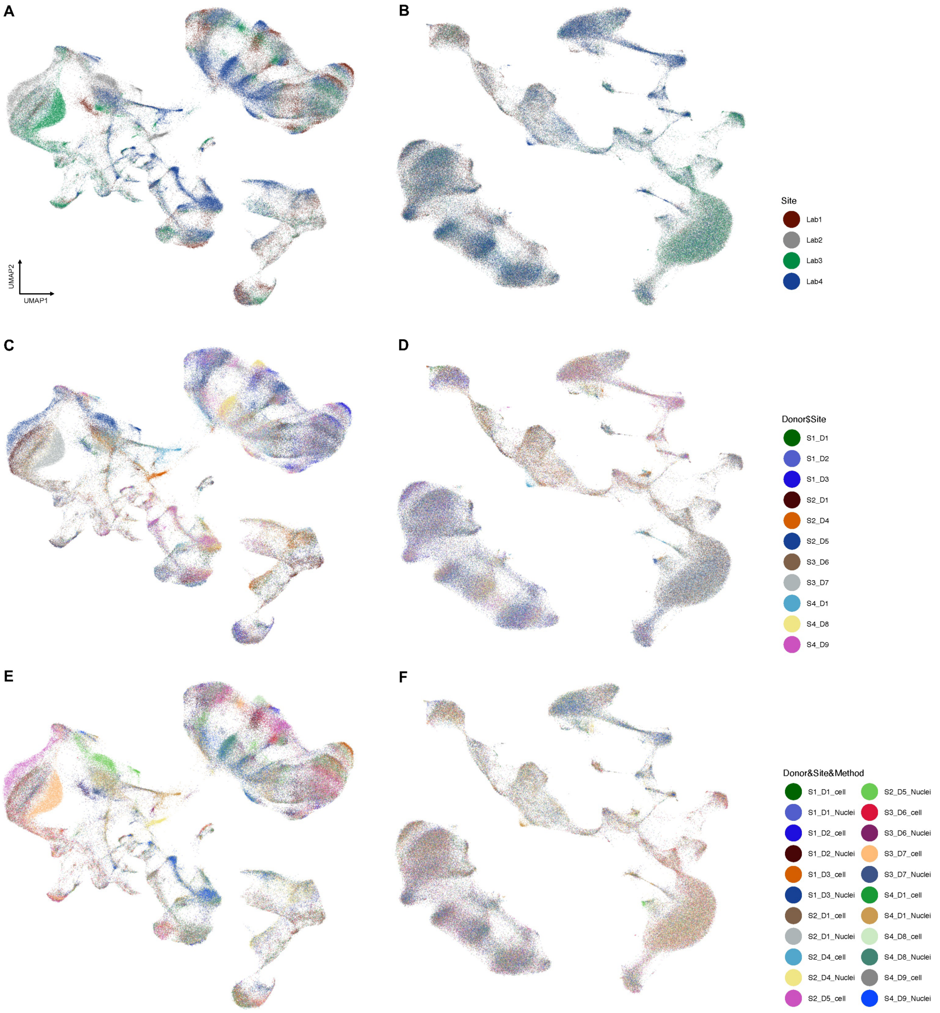


**Figure S11.** UMAP visualisations of the merged datasets before (right) and after batch correction (left), coloured by laboratory(A,B), donors in each site (C,D) and samples (E,F).

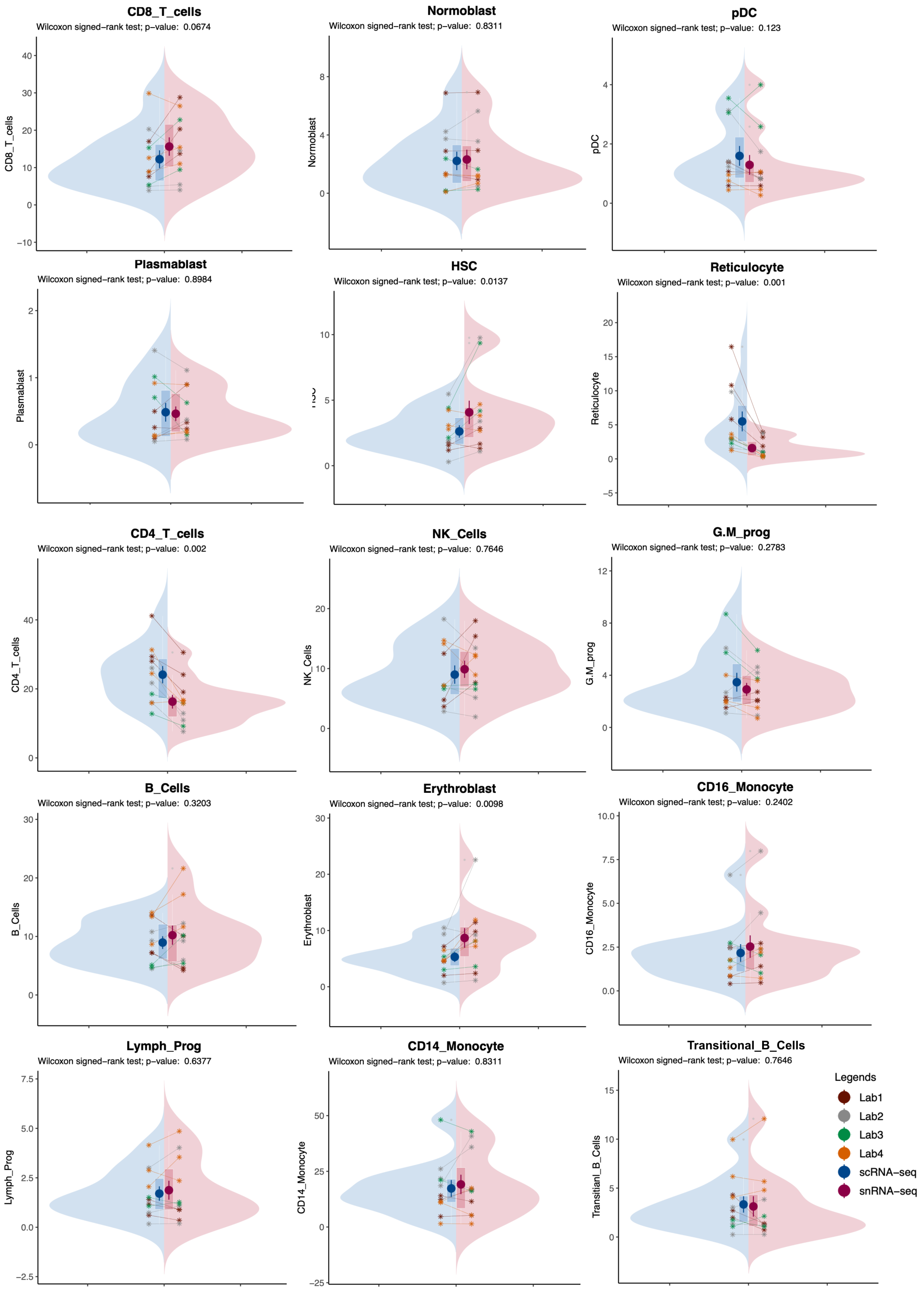


**Figure S12.** Split violin plots showing the proportion of different cell types within each matched sample pair.

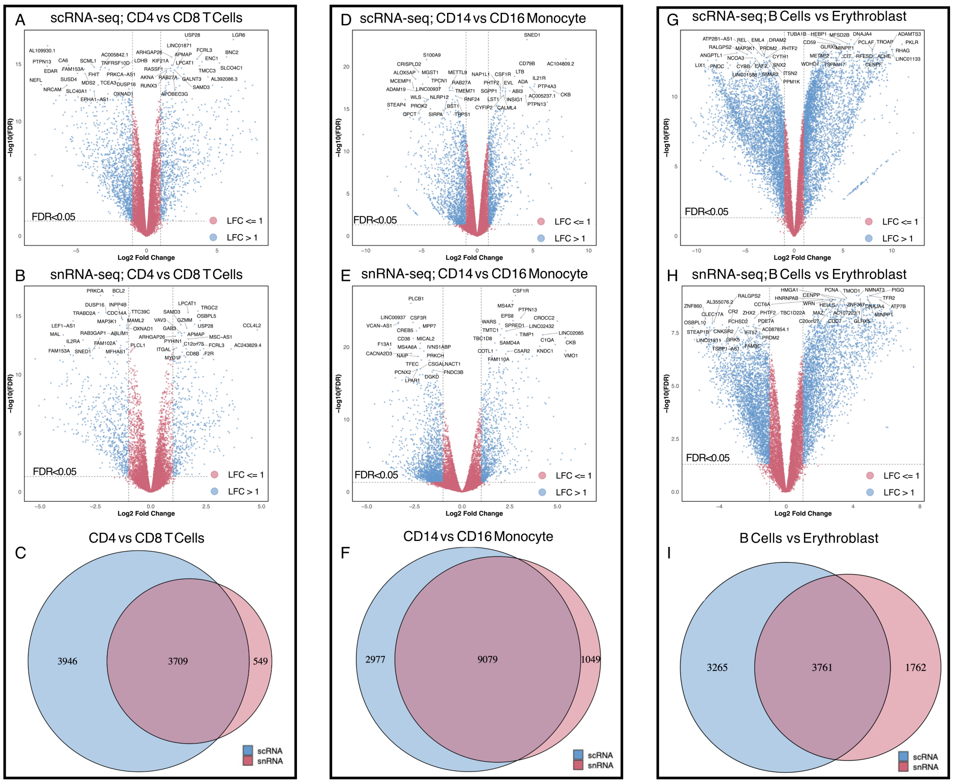


**Figure S13.** Volcano plots showing DEGs from pseudo-bulk analysis of three pairs of cell types within each sequencing methodology (scRNA-seq vs. snRNA-seq), along with a Venn diagram illustrating the overlap of DEGs between the two methodologies. The top 20 upregulated and top 20 downregulated genes in each comparison are labelled.

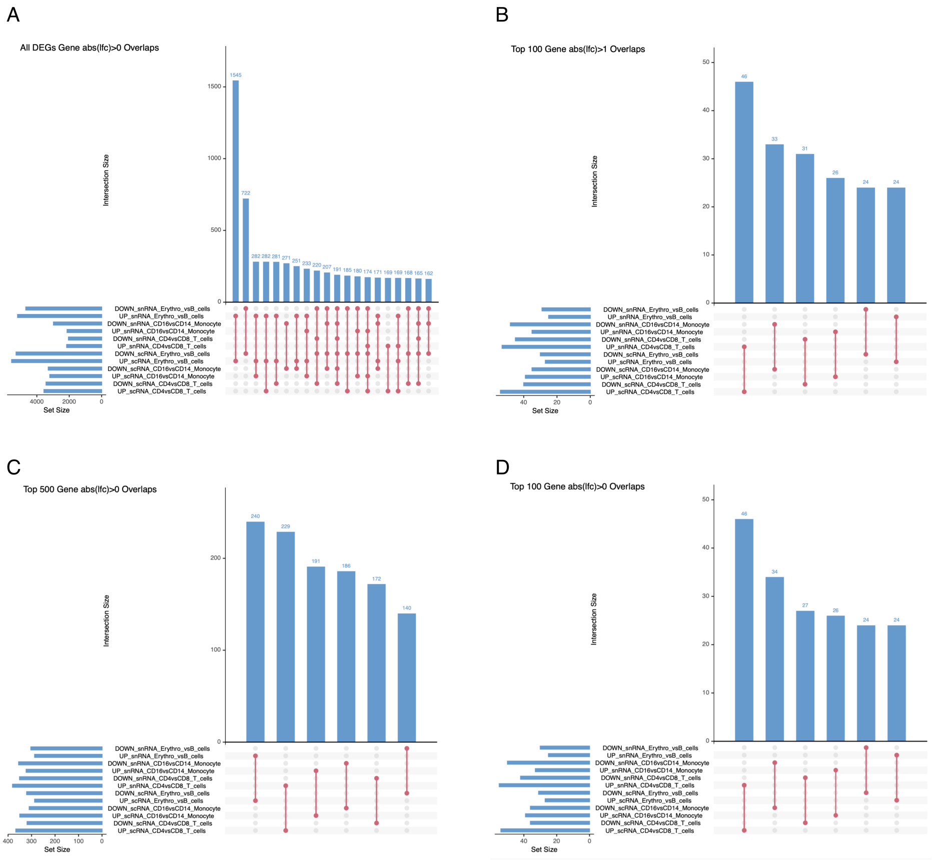


**Figure S14.** Upset plots representing the overlap of DEGs from pseudo-bulk analysis of three pairs of cell types within each sequencing methodology (scRNA-seq vs. snRNA-seq). **A)** All DEGs with FDR $< 0.05$ and absolute log fold change ($|\mathrm{LFC}| > 0$); **B)** Top 100 DEGs with FDR $< 0.05$ and $|\mathrm{LFC}| > 1$, ranked by FDR; **C)** Top 500 DEGs with FDR $< 0.05$ and $|\mathrm{LFC}| > 0$, ranked by FDR; **D)** Top 100 DEGs with FDR $< 0.05$ and $|\mathrm{LFC}| > 0$, ranked by FDR.
